# Phytohormones enhance heavy metal responses in *Euglena gracilis*: evidence from uptake of Ni, Pb and Cd and linkages to hormonomic and metabolomic dynamics

**DOI:** 10.1101/2022.09.22.508751

**Authors:** Hai Ngoc Nguyen, Thien Nguyen Quoc, Duc Huy Dang, Emery R. J. Neil

**Affiliations:** Trent University, Department of Biology, Peterborough, Canada; Trent University, School of the Environment and Chemistry Department, Peterborough, Canada

**Keywords:** *Euglena gracilis*, abscisic acid, cytokinin, cadmium, lead, nickel

## Abstract

Over the last decade, significant effort has been made to understand phytohormonal functions (e.g. cytokinins (CKs) and abscisic acid (ABA)) in metal stress responses of higher plants and algae. Despite the potential for these phytohormones to improve industrial remediation by *Euglena gracilis* (Euglenophyceae), no such roles have been elucidated for this highly adaptive species and its response to heavy metals. This study demonstrates that toxic metals (nickel, lead, cadmium) modify hormonal activity profiles (i.e., CK forms and their concentrations) in *E. gracilis*. Furthermore, exogenous ABA or CK (*t*Z) enabled higher metal uptake efficiency and alleviated metal toxicity through the regulation of endogenous CKs and gibberellins (GAs) levels. These responses suggest that *E. gracilis* regulates multiple phytohormone signals during metal stress acclimation. A deeper approach, using untargeted metabolomic analyses, gave more detailed insight into phytohormone-controlled pathways and associated modified metabolites, which were frequently related to metal accumulation and the physiological acclimation to metal presence. Significant changes in the levels of cellular metabolites, especially those involved in acclimation to metal stress, were under the influence of phytohormones in algal cells. When grown under metal stress conditions, the presence of exogenous ABA or CKs, caused changes in cellular metabolites which included those from: lipid pathways, riboflavin metabolism, the biosynthesis of cofactors/vitamins, and carbohydrate metabolism. Also, bioactive secondary metabolites (e.g., terpenoids, alkaloids, flavonoids, carotenoids) were modified in algal cells treated with phytohormones. Thus, the study gives a detailed view on the regulatory functions of ABA and CKs in algal metal bioremediation strategies, which are attributed to enhanced metal uptake and in the fine-tuning of plant hormone levels during metal stress response. The results can guide efforts to develop efficient, low-cost and environmentally friendly methods for bioremediation.

**Highlights:**

- Metal stress produces phytohormone-specific responses in *Euglena gracilis*.
- Phytohormones (ABA and CK) enhance metal accumulation rates.
- Phytohormone-controlled metal uptake reflects enhanced CK activity profiles.
- Modified hormonal crosstalk is involved in phytohormone-induced metal uptake.
- Metabolomic responses to phytohormones-involve metal stress mitigation compounds.

## 1. Introduction

The mining industry produces one of the largest waste streams in the world. On a global scale, mining annually produces approximately 20-25 billion tons of solid mine waste, and an estimated 5 to 7 billion tons are mine tailings (**Mudd and Boger, 2013**). There is ample evidence suggesting that the continuous and unsustainable impacts of mining industries and other tailing facility failures produce substantial amounts of toxic metal contamination in water supplies, threatening human health and ecosystem balance (**Rai et al., 2019**). Of the numerous toxic heavy metals (HM), nickel (Ni), lead (Pb) and cadmium (Cd) are severely damaging to living organisms, including humans, even at low concentrations. The United States Agency for Toxic Substance and Disease Registry (ASTSDR, 2019) ranked Pb as the second, Cd as seventh and Ni as fifty-eighth of the most hazardous substances **(ASTSDR, 2019)**. These HM are persistent in the environment, and could enter the food chain, posing risks to ecosystems and humans. Therefore, it is of high importance to develop techniques for the removal of these HMs to reduce their environmental toxicity.

There are several physical and chemical approaches for the removal of HMs from the environment, such as, replacement or washing of soil, (co)precipitation, oxidation-reduction, ion exchange, or adsorption **(Cui et al., 2020)**. However, these methods are costly, time- and labor-consuming and more importantly, their success is often limited in natural ecosystems due to the large areas of contaminated land or water (**Cheng et al., 2019**). To overcome this, bioremediation seems to be another efficient, economical and sustainable alternative for the removal of HMs (**Saeed et al., 2021; Tufail et al., 2022**). By using microbes, bioremediation applications can immobilize, remove, and detoxify active heavy metal ions in natural environments (**Liu et al., 2021)**. Many environmental remediation approaches focus on developing microbial bioremediation, which is an *in situ* cost-efficient technology employing bacteria, fungi or algae (**Dar et al., 2019**; **Nguyen et al., 2020**). Microalgae, termed phyco-remediators, have been a strong focus for improvement of microbial systems for HM remediation performance in water (e.g., cadmium, arsenic, chromium, lead, mercury) (**Chugh etal., 2022; Goswami et al., 2022**).

*Euglena gracilis* has attracted attention in both academic and industrial sectors due to its ability to accumulate and neutralize a wide range of heavy metals (**Santiago-Martínez et al., 2015; Khatiwada et al., 2020**). *E. gracilis* is a single cell flagellate eukaryote and photosynthetic organism with a complex evolutionary history. In fact, it may be classified as belonging to the kingdom Protista (**Muchut et al., 2021**) or Algae **(Khatiwada et al., 2020)**. This complex background has contributed to what is now a great metabolic capacity, for producing high levels of diverse bioactive compounds (**Yoshioka et al., 2020**). Evidence indicates that *E. gracilis* has roles in remediating active HM ions by accumulating and transforming them into unharmful ions, or by externally binding and neutralizing them (**Khatiwada et al., 2020**). *E. gracilis* tolerates and accumulates high concentrations of copper (Cu), chromium (Cr), Pb, Cd, mercury (Hg) and zinc (Zn) (**Santiago-Martínez et al., 2015**). To combat the toxic effects of Cd, Cr, Hg and Zn, *E. gracilis* biosynthesizes metal binding compounds such as: cysteine, glutathione, chelating molecules with thiol groups, or phytochelatins. Those compounds effectively reduce toxicity of HMs as a mechanism of cytoplasmic detoxification (**García-García et al., 2020**). These advantages have great potential to be intensified through the optimized application of phytohormones, such as abscisic acid (ABA) and cytokinin (CKs) (**Nguyen et al., 2020; Stirk and Van Staden, 2020**).

The cytokinins (CKs) are phytohormones and key regulators of plant growth and development **(Rashotte, 2020)**. CKs regulate several abiotic stress responses in higher plants (**Nguyen et al., 2020**), including metal stress (**Jiang et al., 2019)**. In micro-algal studies, exogenously applied CK was found to reduce metal absorption and increase Cd, Pb, and Cu tolerance in *Chlorella vulgaris* (**Piotrowska-Niczyporuk et al., 2012**), and applications of CKs improved Pb stress tolerance in *Acutodesmus obliquus* (**Piotrowska-Niczyporuk et al., 2020**). In Euglena, CKs were first bioassayed and putatively identified using gas-liquid chromatography (**Swaminathan and Bock, 1977**). Like CKs, the stress phytohormone known as ABA, is an important regulator of abiotic and heavy metal stress responses in higher plants (**Sytar et al., 2019**). In microalgae, ABA is associated with the action of the 24-epibrassinolide enhancement of Pb stress tolerance in *A. obliquus* and is considered a potential agent for HM bioremediation **(Piotrowska-Niczyporuk et al., 2020**). Discovering additional crosstalk mechanisms among various hormones (ABA, CK, gibberellins (GAs), auxins, salicylic acid (SA), jasmonic acid (JA)) in coordinating growth under metal stress is an important theme in the field of algal bioremediation (**Nguyen et al., 2020**). Hormonal crosstalk is thought to play major roles in HM tolerance via activation of phytochelatin biosynthesis, stimulation of antioxidant apparatus (i.e., carotenoids, ascorbate peroxidase, catalase, superoxide dismutase), or stimulation of protective compounds (e.g., proline, astaxanthin, pigments) (**Piotrowska-Niczyporuk et al., 2020; Talarek-Karwel et al., 2020**).

Our hypothesis was that ABA and CK are influential for the acclimation of algal cells to metal stress. To address this hypothesis, the effects of exogenously applied ABA and CK on *E. gracilis* and their ability to counteract Ni, Pb, and Cd phytotoxicity was examined. Different physiological performance measures were compared among Euglena cultures exposed to the HMs and/or CKs and ABA. These included measures of cell growth and viability, and elemental analysis to determine metal uptake efficiency. Using a targeted hormonomic approach, levels of multiple phytohormones (CK, ABA, GA, IAA, SA) were profiled using high-performance liquid chromatography tandem mass spectrometry (HPLC–MS/MS) to document the occurrence of these phytohormones and their possible crosstalk in metal stress response. The impacts of ABA and CK were more deeply investigated during the accumulation of Ni, Pb and Cd in *E. gracilis* through an untargeted metabolomics approach. The objective was to discover patterns of any key metabolites that are important in determining functional molecular aspects of hormone and HM interactions.

## 2. Materials and methods

Detailed materials and methods are described in supplementary Materials and Methods.

### 2.1 Euglena growth conditions and treatments

Axenic cultures of *Euglena gracilis* Klebs (CPCC 469) were purchased from Canadian phycological culture center, department of Biology, University of Waterloo. Cultures were grown axenically in 50 mL of Modified Acid Medium (MAM) (**Olaueson and Stokes, 1989**), at pH 4.5, in 250 mL plastic flasks. Euglena cultures were maintained on a shaker in an environmental chamber at 24°C. Lighting was supplied by cool-white fluorescent light with an intensity of 97.3 Par under 12/12 light/dark cycle until culture reached 10^6^ cell/mL. Briefly, phytohormones ABA (10^−9^ M) and CK (10^−9^ M *t*Z) were used on their own and in combination with 0.1 mM Pb (as Pb(NO_3_)_2_), 0.05 mM Ni (as NiCl_2_.6H_2_O), or 0.025 mM Cd (as Cd(NO_3_)_2_.4H_2_O). In all testing conditions, there was a control (non-treatment) and nine treatments (Ni; Pb; Cd; ABA; *t*Z; ABA+Ni, ABA+Pb, ABA+Cd, *t*Z+Ni, *t*Z+Pb, *t*Z+Cd). Exogenous phytohormones and metals used in experiments were purchased from OlChemim Ltd (Olomouc, Czech Republic) and Thermo Fisher Scientific, respectively. All supplements were added to 50 mL cultures (using MAM medium) during the early exponential phase (Day 0 with 0.6 × 10^6^ cells/mL) with 4 biological replicates (4 separate 250 mL flasks). Collected cells at the end of treatments (10 days) were washed by deionized water and freeze-dried using a FreeZone freeze dryer (Labconco, Missouri, USA) for subsequent analysis.

### 2.2 Determination of cell growth

For cellular growth characterization of *E. gracilis*, 1 mL samples were collected from treatments after 2, 4, 6, 8 and 10 days of growth and the number of cells determined by Trypan Blue staining **(Khatiwada et al., 2020)**. Enumeration of living Euglena cells was determined by counting cells immobilized with 0.05% HCl, using a 0.1 mm Neubauer hemocytometer (Thermo Fisher Scientific, USA) and a light microscope (Life Technologies AMAFD1000) coupled with EVOS cell imaging systems (Thermo Fisher Scientific, California, USA).

### 2.3 Determination of metal accumulation by Euglena using ICP-MS

At the end of each exposure time, 1mL of sample was filtered using a 0.2 μm syringe tip filter (Canadian Life Science), acidified with double-distilled trace metal grade HNO_3_, and stored at room temperature until elemental analysis. Metal concentrations were determined utilizing a triple quadrupole inductively coupled plasma mass spectrometer (ICP-MS, 8800 Agilent Technologies, Water Quality Centre, Trent University) following the protocol of **Dang et al., (2020**). Briefly, the instrument was operated under He mode to remove polyatomic interferences and Indium was used as an internal standard at a final concentration of 20 µg L^-1^. Analytical recovery was determined using a certified reference material (ES-L1, SCP Science, Quebec, Canada). We also determine metal concentrations in the stock metal solution, blank sample (medium only), Euglena sample (Euglena only), and phytohormone samples (Euglena with phytohormone) for blank and control conditions.

The uptake of metals by *Euglena* was computed using changes in the metal concentration in the test medium during the exposure period expressed as percentage removal: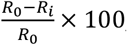, where: R_0_– initial concentration-day 0 and R_i_ -concentration of metal after each exposure period **(Kong et al., 2019; Zada et al., 2021)**.

### 2.4 Cytokinin profiling

#### 2.4.1 Cytokinin extraction

Algal cell pellets (25 mg dry weight) were harvested at the end of the experiment as described above and four replications at final time point (10 days) were used to determine endogenous cytokinin (CK) levels. Extraction, purification, and quantification of CKs were performed as previously described (**Kisiala et al., 2019)**. Samples were extracted in 1 mL of Bieleski solution to which 29 stable isotope-labelled internal standards were added (20 ng of each of the deuterated internal standards, OlChemim Ltd. (Olomouc, Czech Republic)) corresponding to the different forms of CK that were scanned for as listed in previous studies **(Supplementary Table 1) (Kisiala et al., 2019; Aoki et al., 2021; Bean et al.,2021; Palberg et al., 2022)**.

#### 2.4.2 CK Quantification using UHPLC-(ESI+)-HRMS/MS

Endogenous CKs of *E. gracilis* samples were identified and quantified using UHPLC-(ESI+)-HRMS/MS (**Kisiala et al., 2019**). A 25 µl sample volume was injected into the Thermo Ultimate 3000 UHPLC coupled to a Thermo Q-Exactive™ Orbitrap mass spectrometer equipped with a heated electrospray ionization (HESI) source (Thermo Scientific, San Jose, CA, USA). All hormone forms were eluted with a multistep gradient of component A: H_2_O with 0.08% CH_3_CO_2_H mixed with component B: CH_3_CN with 0.08% CH_3_CO_2_H at a flow rate of 0.5 mL/min. The initial conditions of 5% B were held for 0.5 min, increasing linearly to 45% B over 4.5 min followed by an increase to 95% B over 0.1 min; and held at 95% B for 1 min, before returning to initial conditions for 2 min of column re-equilibration, total run time was 8.2 min.

All data were analysed using Thermo Xcalibur (v. 4.1) software (Thermo Scientific, San Jose, CA, USA), to calculate peak areas. Quantification was achieved through isotope dilution analysis based on recovery of ^2^H-labelled internal standards. Blank media samples (control, MAM solution) were processed and analysed using the same methodologies shown no background levels of CKs.

### 2.5 Hormonomics

ABA, indoleacetic acid (IAA), SA, and selected Gibberellins (GA_1_, GA_3_, GA_4_, GA_7_) were scanned for in 25 mg dry weight *E. gracilis* pellet samples collected after 10 days of treatment. Extraction, purification, and quantification of phytohormones was performed as described by **Cao et al (2020)**.

All samples were analyzed in negative ion mode using a HPLC–ESI–MS/MS (QTrap5500 triple quadrupole mass spectrometer (Sciex Applied Biosystems, Massachusetts, United States) connected to a Shimadzu LC10ADvp equipped with PE200 autosampler. A 20 uL aliquot was injected onto a reverse phase C18 column (Kinetex 2.6u C18 100 A, 2.1 × 50 mm; Phenomenex). Phytohormones (ABA, IAA, GA_1_, GA_3_, GA_4_, GA_7_ and SA) were eluted using component A: H_2_O with 0.5% formic acid (FA) and component B: 0.5% FA in ACN (v/v), at a flow rate of 0.5 mL min^−1^. Phytohormones were analyzed in negative ionization mode and the programmed step gradient was the same as **Cao et al., (2020)**.

### 2.6 High resolution Orbitrap mass spectrometry-based metabolomics

For the metabolomics extractions, 25 mg dry weight pellet samples were collected from experiments described above and subsequently extracted following **Renaud et al. (2017)**. Labeled standards (10 ng of [^2^H_7_]BA and 10 ng of [^2^H_6_]ABA) were added as lock masses and to monitor retention times across samples. Full scan analyses were performed following **Renaud et al. (2017)** with modifications, using UHPLC-MS. The samples were passed into the mass spectrometer from a Thermo Ultimate 3000 UHPLC system (FisherScientific, Ottawa, Canada). For full scan analysis, each sample was run in negative ion mode over the mass range of m/z 80−600, at 35,000 resolutions, with automatic gain control (AGC) target of 3 × 10^6^, and maximum injection time (IT) of 128 ms. For metabolite selection, data files (as Thermo.RAW) were converted to mzXML file by ProteoWizard (Version 2.1) and uploaded to XCMS Online (https://xcmsonline.scripps.edu) to query significant differences among the metabolite profiles **(Papaioannou et al., 2021; Rodríguez-Moro et al., 2022)**. To ensure the sensitivity, reliability and validity of metabolite profiling analyses, specific internal standards were monitored in either negative (ABA) or positive modes (Benzylaminopurine), and samples were analyzed at three dilution levels (1x, 10x and 100x dilution factor) (**Aoki et al., 2019; Palberg et al., 2022)**. The optimization tests showed that undiluted samples in negative mode were the most robust and so these parameters were selected for subsequent analysis. High abundance metabolites (fold change ±1.5, p-value < 0.05) in the *E. gracilis* are summarized in **Figure 4**. Examples of highly selective extracted ion chromatograms (EIC of significantly up/down regulated metabolites are shown in **Supplementary Figures 3.1-3.9)**.

### Metabolite Identifications

All putative feature identifications were performed to a Level 3 confidence assignment – according to a scale set by the Chemical Analysis Working Group (CAWG) of the Metabolomics Standards Initiative (MSI) and as modified by **Schrimpe-Rutledge, (2016)**. Accordingly, accurate masses of features were queried against databases to produce tentative structures and identifications.

MetaboQuest (http://tools.omicscraft.com/MetaboQuest/) was used as a single search tool and was queried with the m/z value of each unfragmented feature as a positive ion, with [M+H] and the [M+Na], [M+ACN] adducts specified. Functional annotations of the proposed metabolites were derived using MetaboQuest (http://tools.omicscraft.com/MetaboQuest/), PubChem (https://pubchem.ncbi.nlm.nih.gov/) and KEGG **(Cheng et al., 2022; Rodríguez-Moro et al., 2022)**.

### Statistical analyses

Statistical analyses were performed with SPSS Software (IBM, version 28.0) on algal physiology and biochemistry data using a one-way analysis of variance (ANOVA) followed by Dunnett’s post-hoc test to test for significant effects of treatments when compared to the respective controls. Significant differences (P-< 0.05) for the Dunnett’s test determine significant effects of treatments as compared to a control (non-treated group). Data points and error bars reflect means ± standard errors of 4 biological replicates for phenotyping and phytohormone data and at least three biological replicates for metal uptake analyses.

## 3. Results

### 3.1 The effect of phytohormones on heavy metal accumulation

The capacity of Euglena to accumulate metals was determined using ICP-MS at different culture time points (2, 4, 6, 8 and 10 days) (**Figure 1**). There was a steady bioaccumulation of Ni (percentage-%) throughout the experiments (day 2-day 10) and it peaked at day 6 (76.8%) and trailed off to a minimum of 70.1% by day 10 **(Figure 1a)**. As a standalone treatment, Pb accumulated gradually from day 2 to a maximum of 13.3% at day 4 (**Figure 1b**). Afterward, the Pb content decreased to a minimum of 0.85% by day 10. The maximum for Cd accumulation by the Euglena was more than 52% by the end of treatment with only a small change from day 4 to day 10 (**Figure 1c**).

**Figure 1.**
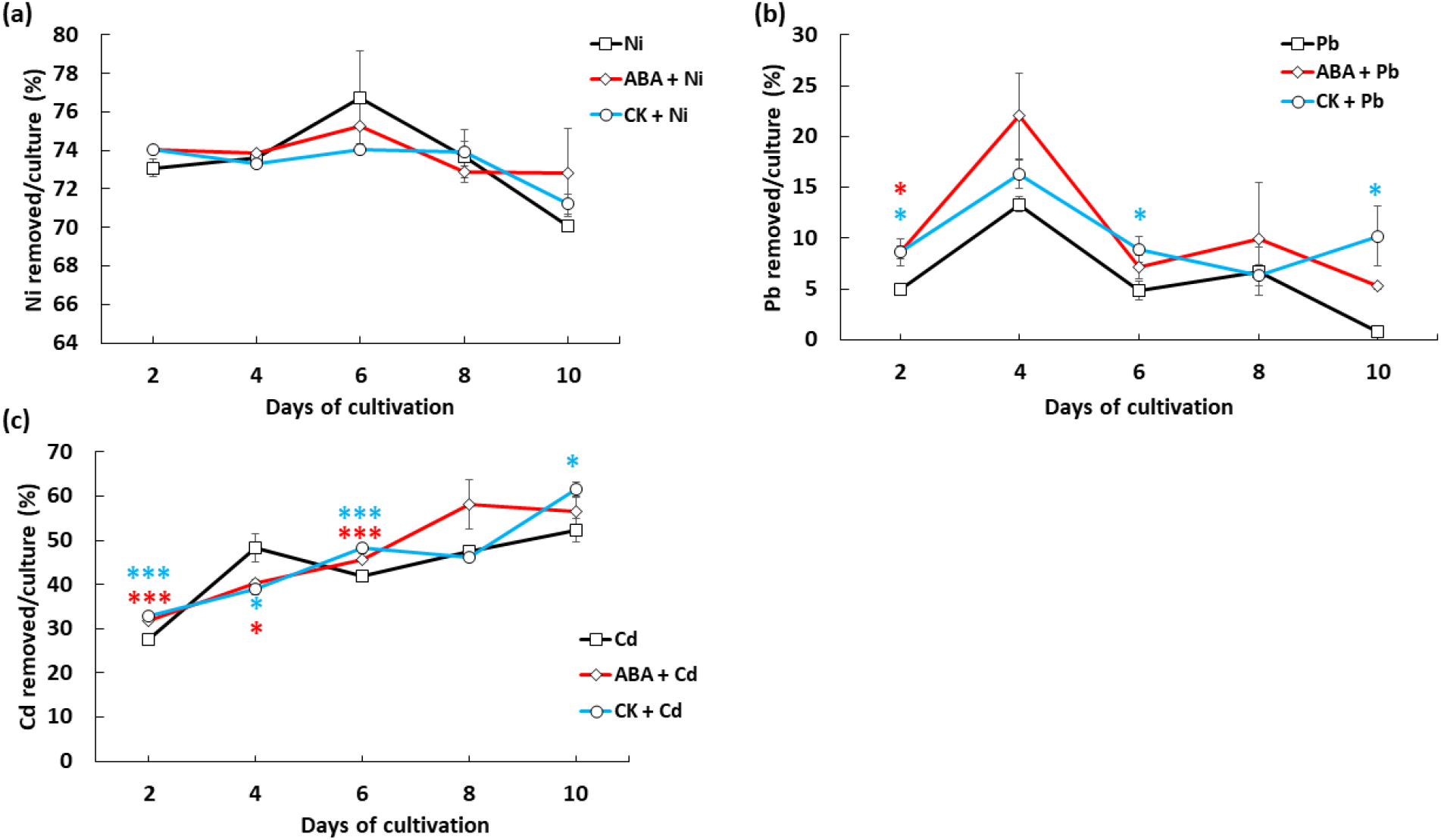
Phytohormones enhance metal uptake efficiency in Euglena. The effect of CK and ABA on removal capacity (%) of *E. gracilis* for three heavy metals Nickel (0.5 mM) (a), (Lead (0.1 mM) (b), Cadmium (0.025 mM) (c). Indium (20 µg L^-1^) was used as an internal standard for ICP-MS analysis. The removal of metals by *Euglena* was calculated using changes in the metal concentration in the test medium during the exposure period expressed as percentage removal: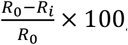, where: R_0_- initial concentration-day 0 and R_i_ - concentration of metal after each exposure period. For validation, metal concentrations in the stock metal solution, blank sample (medium only), Euglena sample (Euglena only), and phytohormones sample (Euglena with phytohormone) were determined to blank and control conditions. The data shown are for time-course of the (a) lead, (b) nickel and (c) cadmium removal at 2, 4, 6, 8, 10 days, respectively. Data are the means of at least three independent biological replications ± SD **(Supplementary Table 4)**. The difference in removal capacity between the treatments with phytohormone (red for ABA, blue for CK) was significantly different (*) accepted when marked by * (* P < 0.05; **P<0.01, ***P<0.001 versus control using Dunnett’s test). The lack of * at indicates there were no significant differences.

When combined with heavy metals, a cytokinin, *trans*-Zeatin (*t*Z), induced a higher increase in Cd uptake; Increases under Cd stress, were as follows at day 2 (5.4%, p < 0.001), day 6 (6.2% p < 0.001), and day 10 (9.2%, p < 0.05) **(Figure 1c)**. trans-Zeatin with Pb treatment also increased Pb uptake at day 2 (1.7 fold, p < 0.05), day 6 (1.8 fold, p < 0.05), and day 10 (12.0 fold, p < 0.05) (**Figure 1b**). ABA supplementation in combination with Pb consistently increased Pb uptake at days 2, 4, 6, and 8; whereby It was significantly greater than controls at day 2 (1.7 fold, p < 0.05), and trending higher at day 6 (1.47 fold, ns), and day 8 (1.48 fold, ns) **(Figure 1b)**. A stimulatory effect of ABA on metal uptake was also observed in case of Cd on days 2, 6, 8, and 10 (16.0% (p < 0.03)), 9.0% (p < 0.013), 22.1% (ns), and 8.1% (ns), respectively). Though not significant, both *t*Z and ABA, when co-treated with Ni, appeared to improve Ni uptake efficiency at day 2 (0.98%, p=0.055 for ABA; 0.97%, p=0.057 for *t*Z, **Figure 1a**). Neither *t*Z nor ABA had any significant effect of Ni uptake at any other time point.

Metal toxicity, in the form of culture growth inhibition, for Ni, Pb and Cd was determined for *E. gracilis* cultures and compared to cell growth controls in the absence of heavy metals **(supplementary figures 1 and 2)**. There was a decrease in cell number by 29.7% (p < 0.001, Dunnett’s test), 46.9% (p < 0.001) and 34.4% (p < 0.001) in response to Ni, Pb and Cd, respectively, by the 10th day of the treatment **(supplementary figure 1)**. There were no significant mitigative effects (P < 0.05) of ABA or CK on final cell numbers **(supplementary figure 1)**.

**Figure 2.**
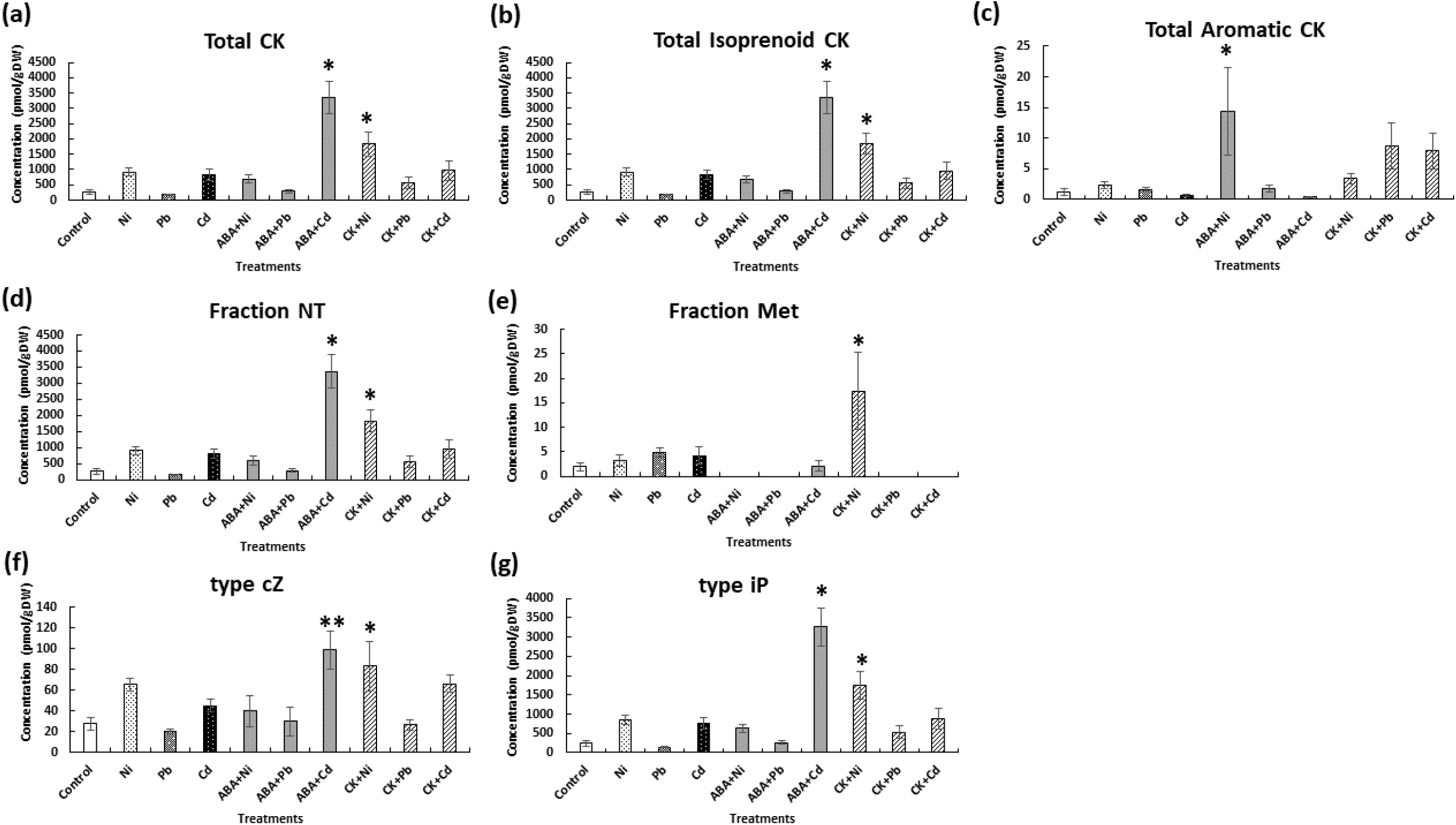
The effect of exogenous ABA and CK (*t*Z) on the level of different forms of endogenous CKs in *E. gracilis* cells exposed to nickel, lead, and cadmium on the 10^th^ day of cultivation in relation to control. Concentration of CK (a) total CKs expressed, (b) Isoprenoid CKs (Free bases+ Ribosides+ Nucleotides+ Methylthiols), (c) aromatic CK (oT+BAP), (d) Nucleotides (NT) (*t*ZNT+ *c*ZNT+ DHZNT+ iPNT), (e) Methylthiols (Met) (2MeSZ+2MeSZR + MeSiPA), (f) type *cis*-zeatin (*c*Z + *c*ZR + *c*ZNT), and (g) type isopentyladenine (iP + iPR + iPNT) as means of four independent biological replications (n= 4) ± SE. Asterisks denote significant differences among treatments in a phytohormone level compared to the control culture (ANOVA; Dunnett’s test, P < 0.05). The lack of asterisks indicates there were no significant differences.

### 3.2 The effect of exogenous ABA and *t*Z treatment on endogenous hormonomics

#### Endogenous cytokinin profiles

The endogenous CK profiles in *E. gracilis* cells under different metal/phytohormone treatments are shown in **Figure 2** and **Supplementary Table 2**. Metal treatments alone caused some notable increases in cell CK contents. Cd treatment significantly induced isoprenoid CKs while none of the metal-alone treatments affected aromatic CKs **(Figure 2b and 2c)**. Both Ni and Cd caused over a 3-fold increase in total CK **(Figure 2a)** and in the case of Cd the difference was significant (P < 0.05). This same pattern was observed for nucleotides CK and iP-type-CK **(Figure 2d and 2g)**. For 2-methylthiols-CKs all three metals resulted in slight but non-significant increases **(Figure 2e)**.

When metal treatments were combined with phytohormone treatments, some dramatic and significant increases occurred in two of the HM/phytohormone combinations. Notably both ABA+Cd and *t*Z+Ni caused large and significant accumulations in: total CK, total isoprenoid-CK, NT-type-CK, *c*-Z type-CK, and iP-type-CK **(Figure 2a, b, d, f, g). Figure 2e** shows 2-methylthiols-CK significantly increased when treated with CK+Ni, but not for the ABA-Cd treatment. The combined treatment of ABA+Cd and CK+Ni was found to enhance the level of cZ-type-CKs (**Figure 2f)**. Another remarkable significant increase in CK was that of an 11.2-fold increase in total-aromatic CKs when treated with ABA+Ni (**Figure 2c)**.

The application of exogenous ABA to the cultures treated with Ni increased the endogenous total CK level as compared to untreated controls (2.5 fold) **(Figure 2a)**. Total CK content was stimulated in cells treated with ABA + Cd by 12.4 fold (p < 0.0001) as compared to the control and by 4.0 fold as compared with Cd metal only (p < 0.05) **(Figure 2a)**. The individual CKs that contributed to the changes seen in the CK groups of Figure 3 are detailed in **Supplementary Table 2**.

**Figure 3.**
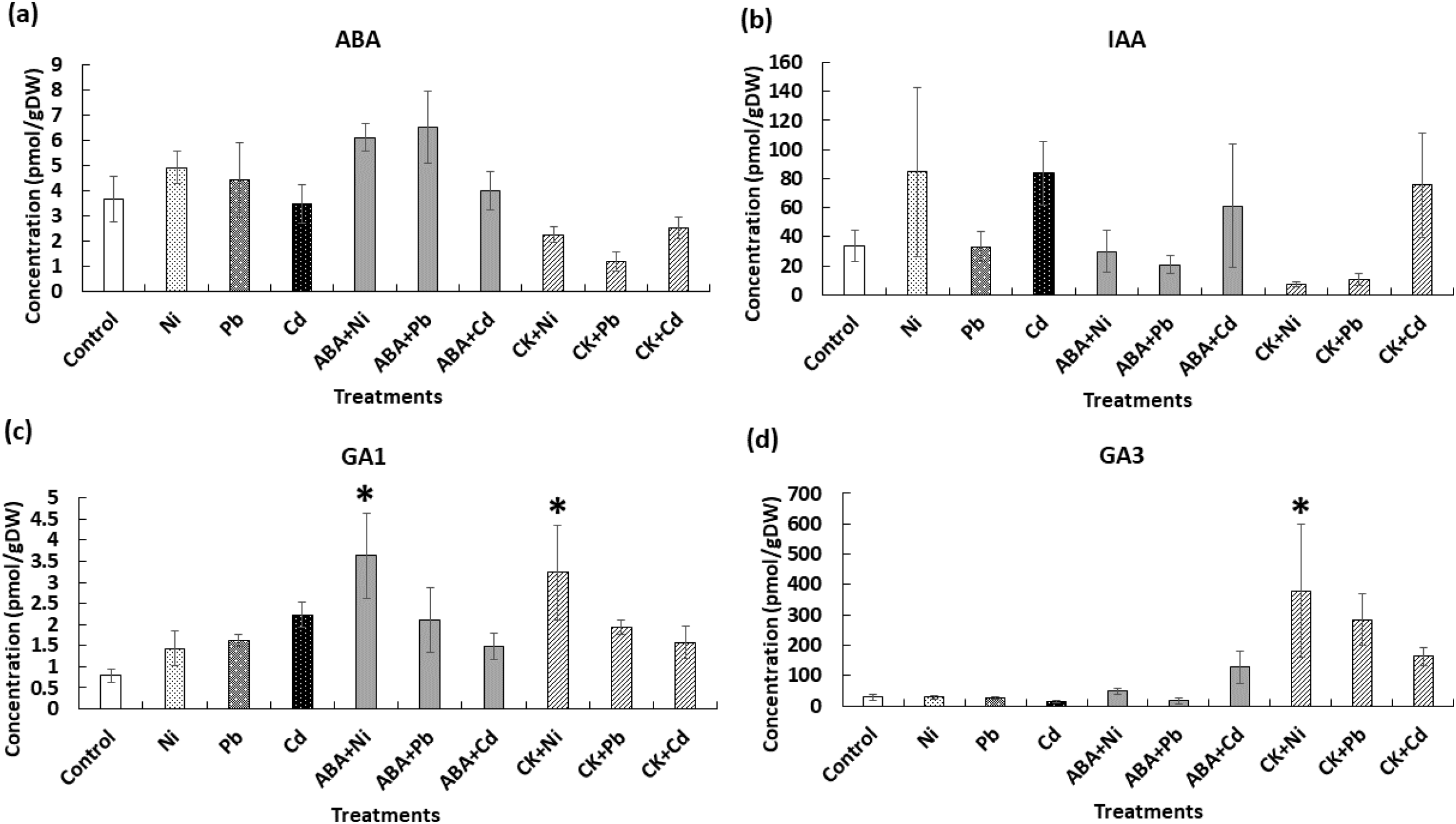
The effect of exogenous ABA and CK (*t*Z) on the level of endogenous (a) abscisic acid (ABA), (b) IAA, (c) (GA_1_), and (d) (GA_3_) in *E. gracilis* cells exposed to nickel, lead, and cadmium, on the 10^th^ day of cultivation in relation to control. Data are the means of four independent biological replications ± SE. Asterisks denote significant differences among treatments in a phytohormone level (ANOVA; Dunnett’s post hoc, P < 0.05). The lack of asterisks at indicates there were no significant differences.

**Figure 4.**
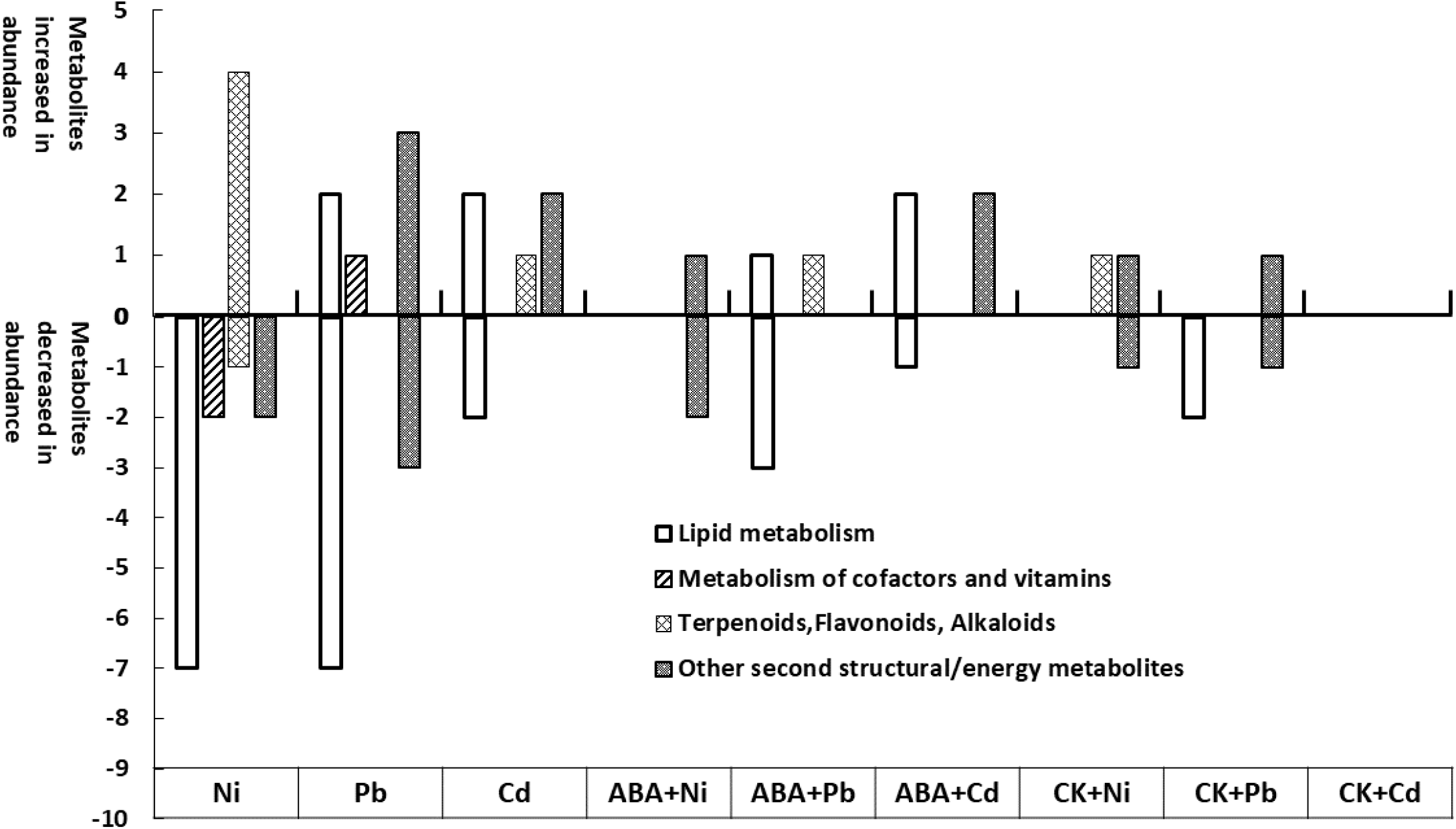
High abundance metabolites (fold change ±1.5, p-value < 0.05) in the *E. gracilis* according to the High Resolution Orbitrap Mass Spectrometry-based metabolomics analysis (three independent biological replications). The metabolites in each category, obtained after heavy metal exposure are represented by bars indicated with numbers of metabolites. Metabolite selection was performed using XCMS Online (https://xcmsonline.scripps.edu). Functional annotation was derived using MetaboQuest and KEGG (https://www.genome.jp/kegg/; http://tools.omicscraft.com/MetaboQuest/). High abundance metabolites in *E. gracilis* are listed in **Table 1** and **Supplementary figure 3**, respectively.

Total isoprenoid CKs (free bases+ ribosides+ nucleotides+ 2-methylthiols) that were detected in *E. gracilis* are shown in **Figure 2b**. Nickel exposure was associated with a strong induction of isoprenoid CK content (3.4 fold higher than control). Notably, Cd treatment had a significant effect on isoprenoid CK (2.5 fold higher, p < 0.001). Exogenous *t*Z induced changes in endogenous isoprenoid contents in algal cultures exposed to Ni (6.8 fold, p < 0.01,), Pb (2.1 fold, ns) and Cd stress (3.6 fold, ns). The level of total endogenous aromatic CKs (oT+BAP) was unchanged from the level of control cultures in response to three metals (**Figure 2c**). ABA applied together with Ni induced a large increase in aromatic CK content (12.5 fold, p < 0.05).

Exogenous ABA and *t*Z induced changes in endogenous contents of inactive CK conjugates, the CK nucleotides (*t*ZNT+ *c*ZNT+ DHZNT+ iPNT), in algal cultures exposed to heavy metal stress (**Figure 2d and 2e**). The levels of NTs increased under ABA+Ni and ABA + Cd (p < 0.0001) treatment by 2.6 and 12.5 fold, respectively. The supplementation of *t*Z to the cultures exposed to each of the three metals increased production of CK NTs as compared to the control (non-treatment) (by 6.8, 2.0 and 3.6 fold (ns) for Ni, Pb and Cd, respectively). The highest increase in *c*Z type CKs (*c*ZNT) was observed in cells exposed to ABA + Cd (3.5-fold increase over the control, ns) and *t*Z + Ni (3.0 fold higher than control, p < 0.05) (**Figure 2f**). Following the results of **Aoki et al., (2021)**, we selected an inorganic media (MAM) known for its negligible background CK levels and we confirmed this was the case for the Euglena samples grown in MAM.

#### Endogenous ABA, IAA, GAs, and SA profiles

Analysis of *E. gracilis* cells indicated that none of any of the measured hormones (ABA, GAs (GA_1_, GA_3_) or IAA) showed any significant change when exposed to any of the three HMs alone. Three of the hormones were never detected (SA, GA_4_, GA_7_).

When HM were combined with *t*Z or ABA treatment, some notable patterns emerged. Regarding **ABA (Figure 3a)** all the CK+HM treatments caused a decrease in ABA levels, although these were not significantly different. For IAA **(Figure 3b)** any treatment containing Cd, caused an increase in cellular IAA (not significant). Two GAs were detected, GA_1_ and GA_3_, and these are both active forms. For GA_1_ both Ni and hormone treatments (ABA+Ni and *t*Z+Ni) caused significantly greater GA_1_ levels (4.6 fold for ABA+Ni, p < 0.05; 4.1 fold for *t*Z+Ni, p < 0.05; **Figure 3c**). Likewise, there was a significant increase in GA_3_ accumulation after *t*Z+Ni treatment. In fact, all the CK+HM treatments caused GA_3_ increases, though only the CK+Ni was significant.

### 3.3 The effect of exogenous phytohormones on intracellular metabolite profiles

To investigate the regulatory function of ABA and CK with a view of gaining insight into the molecular aspects of hormone-mediated heavy metal stress responses, comparative metabolome profiling was performed with HMs in combination with ABA or CK (Phytohormone+metal vs metal alone). Only highly significant features from extracted ion chromatograms selection were used for metabolite identification and functional annotation (whereby only the most highly abundant metabolites were categorized in **Table 1a-1i** and **figure 4)**. Metabolite profiles of *E. gracilis* were substantially changed after exposure to metal when compared to the unexposed cultures (comparisons between control with Ni treatment are shown in **Table 1a**, control with Pb treatment shown in **Table 1b**, and control with Cd treatment shown **Table 1c**). Also, hierarchical clustering heatmap analyses demonstrated there were clear differences among the three metal treatment groups (Ni or Pb or Cd) as compared to phytohormone combined metal treatments (ABA+metal or CK+metal) (data not shown). Comparisons of metabolite profiles were as follows: Ni treatment and ABA+Ni (**Table 1d)**; Pb treatment and ABA+Pb s(**Table 1e)**; Cd treatment and ABA+Cd (**Table 1f)**; Ni treatment and CK+Ni (**Table 1g)**; Pb treatment and CK+Pb (**Table 1h)**; and Cd treatment and CK+Cd (**Table 1i)**. The proposed annotation of metabolites that changed in relative abundance in the *E. gracilis* cell pellets revealed major groupings of metabolites involved in the cellular responses to heavy metals **(Table 1d-1i**). Some of the proposed key features are discussed below based on their accurate mass m/z putative identifications.

**Table 1.**
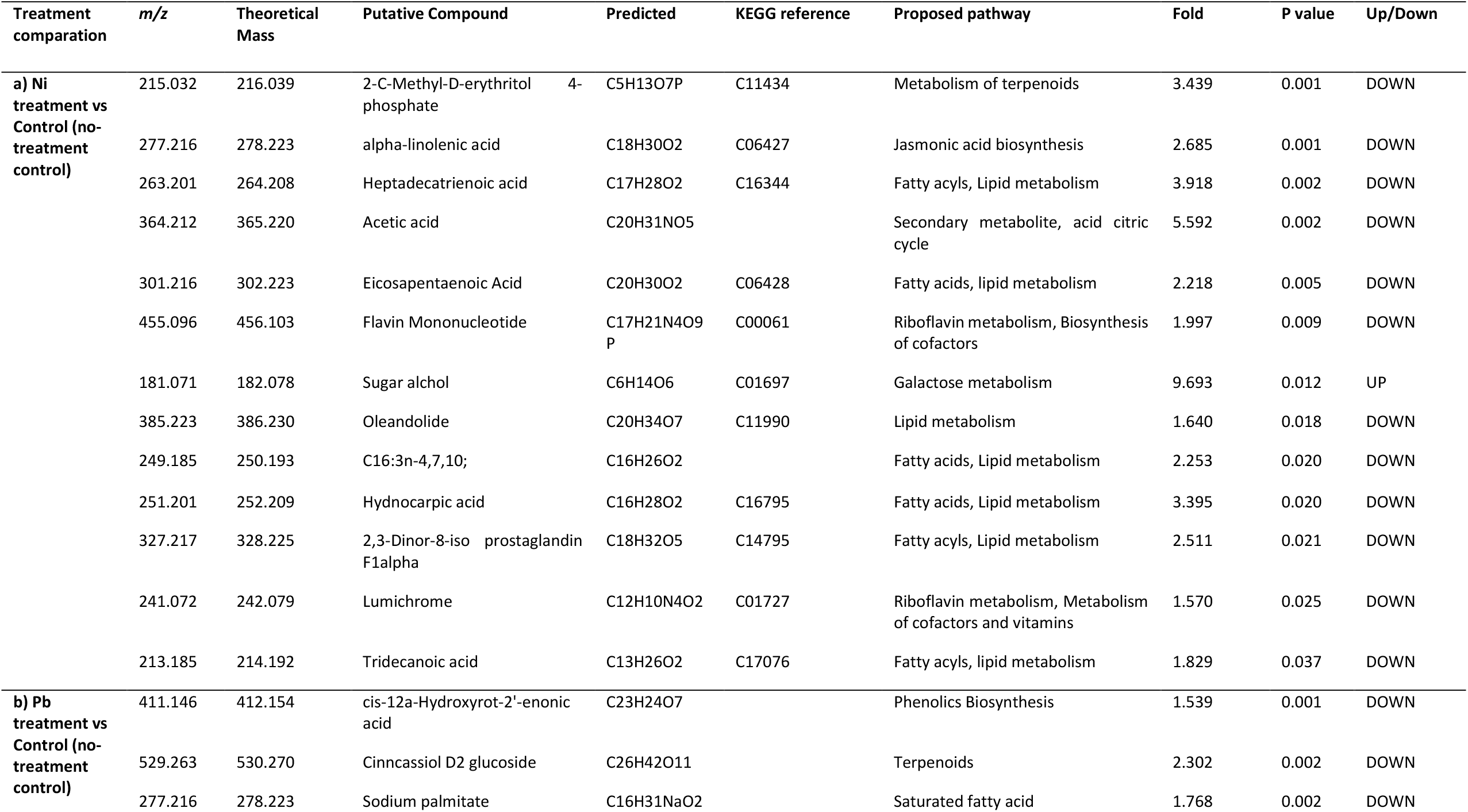
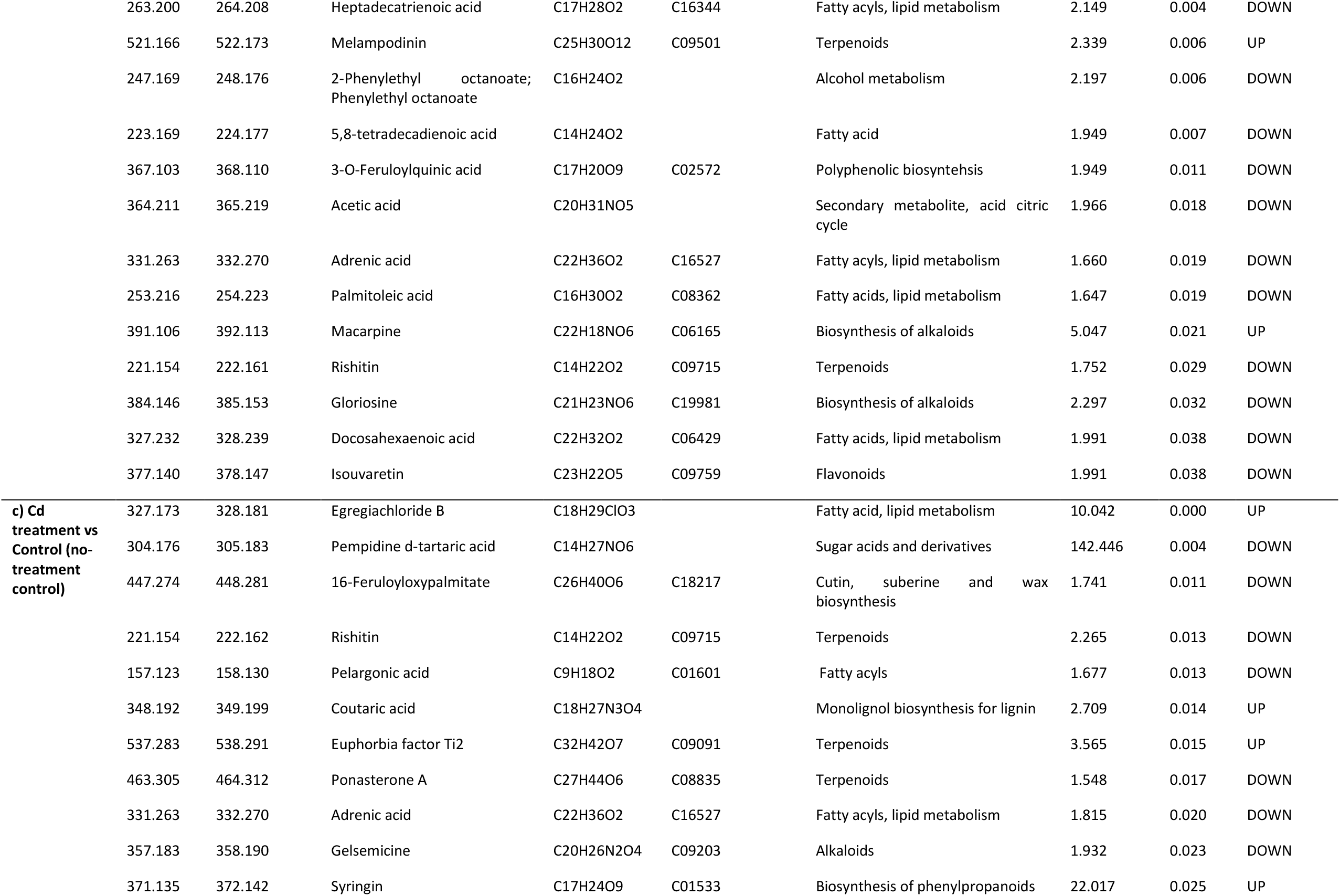
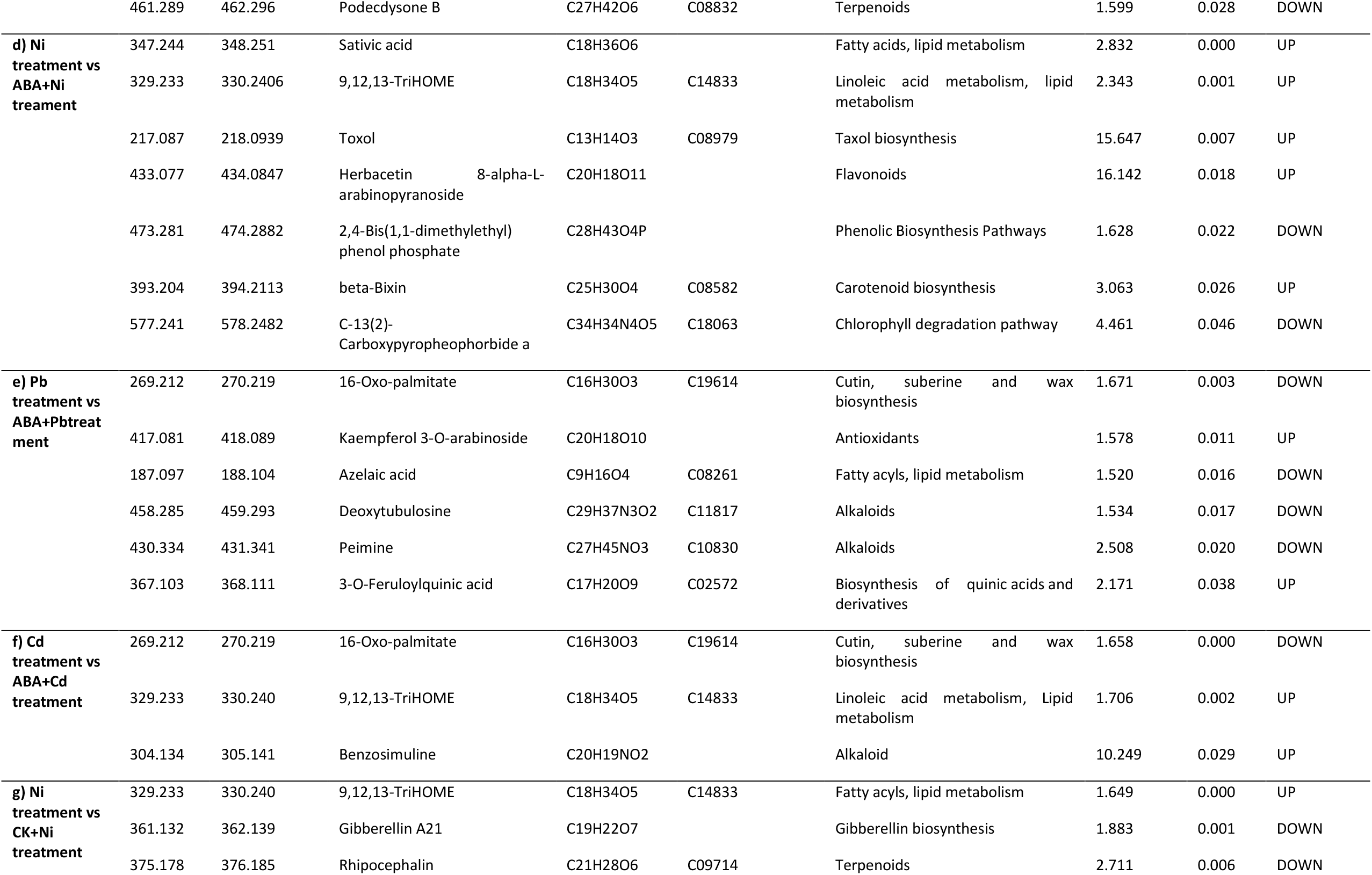
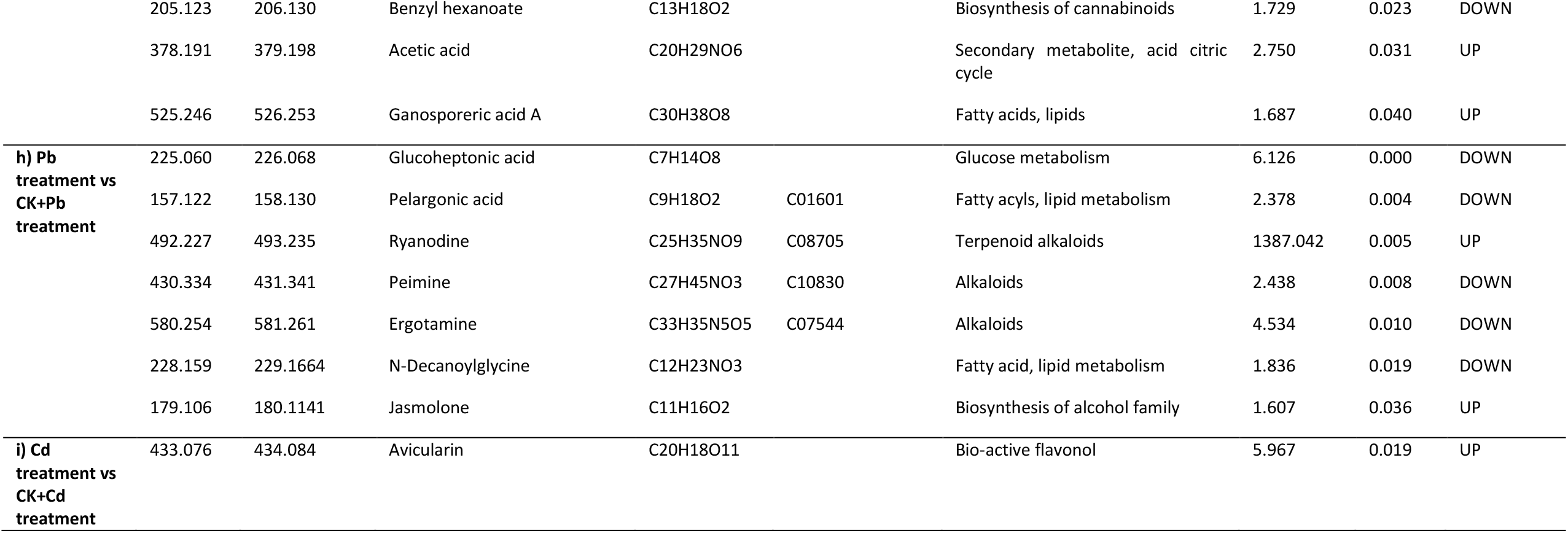
Annotated *E. Gracilis* metabolites analyzed by High Resolution Orbitrap Mass Spectrometry-based metabolomics analysis (three independent biological replications). Metabolites which levels significantly changed in *E. Gracilis* cells exposed to metal and ABA or CK (*t*Z) at harvest time point (10^th^ day of cultivation). Metabolite selection was performed using XCMS Online (https://xcmsonline.scripps.edu). Functional annotation was derived using MetaboQuest and KEGG (https://www.genome.jp/kegg/; http://tools.omicscraft.com/MetaboQuest/).

#### Metabolite profiles of *E. gracilis* under metal exposure

The *E. gracilis* cells exposed to Ni, Pb and Cd showed changed abundance of a variety of compound groups, including: those involved in metabolism of terpenoids, fatty acids and lipids, cofactor/vitamin, amino acids, phenolics, and active secondary metabolites. Exposure to Ni caused a decrease in the production of riboflavin metabolism compounds, such as flavin mononucleotide (1.99 fold) and lumichrome (**Table 1a**). Lead exposure doubled melampodinin (terpenoids) level in *E. gracilis* cell pellets as compared to the control (**Table 1b**). Also, some of the other Pb-downregulated compounds belong to terpenoid (cinncassiol D2 glucoside, rishitin), and flavonoid (isouvaretin) classes (**Table 1c**). The large decrease in the relative abundance of the potentially active secondary metabolites including alkaloids (macarpine, gloriosine, gelsemicine), was observed in algal cultures treated with Pb and Cd, (**Table 1b and 1c**).

#### Metal Stress Metabolites modified by ABA

##### Lipids and related compounds

For *E. gracilis* cells growing in the presence of Ni, ABA induced an increased the content of fatty acids such as sativic acid (2.83 fold) and a linoleic acid precursor (KEGG: C14833, 2.343 fold) (**Table 1d**). Generally, ABA caused significantly reduced levels in algal lipids in Pb-treated cultures, when compared to Pb-alone (1.67 fold of 16-Oxo-palmitate, 1.52 fold azelaic acid, and 2.01 fold of N-decanoylglycine). The relative increases in the levels in the compounds of linoleic acid metabolism (1.71 fold) were observed in algal cultures treated with ABA and Cd.

##### Secondary Metabolites and Pigments

Co-application of ABA with Ni increased the level of secondary metabolites. Examples of these included: flavonoids (herbacetin 8-alpha-L-arabinopyranoside, 16.14 folds) and carotenoids (beta-bixin, 3.06 fold) (**Table 1d**). Likewise, ABA up-regulated benzosimuline under Cd stress (10.25 fold).

#### Metal Stress Metabolites modified by Cytokinins

Accumulations of metal binding compounds were observed in the three metal treatments when supplemented with *t*Z (*t*Z+Ni, *t*Z+Pb, *t*Z+Cd) compared to the cultures treated with metals alone (**Table 1g, 1h and 1i**, P<0.05).

The active CK, *t*Z, in combination with Ni, down-regulated rhipocephalin (terpenoids, 2.71 fold) (**Table 1g**). The addition of *t*Z + Pb-treated cultures modified the synthesis of fatty acids, by downregulating pelargonic acid (2.38 fold), and N-Decanoylglycine (1.84 fold) and upregulating sativic acid (1.56 fold) (**Table 1h**). When comparing the mixture of *t*Z + Pb to Pb only, there was a lowering of carbohydrate metabolism (as indicated by glucoheptonic acid by 6.13 fold (**Table 1h**). Application of exogenous *t*Z stimulated the synthesis of bioactive flavonol (Avicularin, 5.97 fold) in *E. gracilis* exposed to Cd stress (**Table 1i**).

## 4. Discussion

Microalgae have sophisticated mechanisms to physiologically acclimate and adapt to heavy metal stress. One strategy is to accumulate excess, toxic metals in the cells as less active bound forms or in subcellular compartments such as vacuoles, chloroplasts, and mitochondria **(Moreno-Sánchez et al., 2017)**. To this end, they may sequester metals using thiol-compounds (Cys, gammaglutamyl cysteine (γEC), GSH and Poly-GSH) and mitigate damage through activation of antioxidants (superoxide dismutase-SOD, catalase-CAT, peroxidase-POD), or produce global stress response/defense compounds (i.e., transcriptional factors, cellular metal transporters, mitogen-activated protein kinases-MAPKs, chaperones) (**Khatiwada et al., 2020; Hernández-Garnica et al., 2021)**. These strategies can be regulated by phytohormones such as cytokinins (CKs) and abscisic acid (ABA) **(Piotrowska-Niczyporuk et al., 2020; Talarek-Karwel et al., 2020)**. Recent studies have started to suggest that CKs and ABA can regulate cellular redox balance, nutrients and other phytohormones in HM stress responses **(Piotrowska-Niczyporuk et al., 2012, 2018, 2020; Talarek-Karwel et al., 2020**). Given this scenario, our study has detailed an effect of CK- and ABA-induced enhancements of metal uptake, and underlying mechanisms of hormone cross-talk and metal acclimation pathways. These insights can be used for future microalga biotechnological discoveries including genetic modification of algae for improved bioremediation of heavy metal pollution.

### 4.1 Phytohormones enhance metal uptake efficiency in Euglena

The results of phenotyping and metal uptake efficiency indicated that exogenous ABA and a CK (*t*Z) applied with two heavy metals (Pb or Cd) have strong positive influences on the capacity of metal accumulation in *E. gracilis*, but they do not significantly mitigate cell culture growth rate inhibition caused by metals **(Figures 1 and Supplementary Figure 2)**. Even though exogenous hormone applications do not improve cell population growth, they do allow for better maintaining of viability for the existing cells **(Supplementary Figure 2)**. Unlike for the cases of Pb and Cd, exogenously applied ABA or *t*Z did not improve metal uptake in algal cells treated with Ni **(Figure 1)**. Pb, Ni, and Cd can have toxic effects on green microalga *E. gracilis*, reducing the cell number growth in the cultures **(García-García et al., 2018; Khatiwada et al., 2020)**. Algal growth suppression may be associated with hormone balance disruptions that can involve CKs, ABA, auxin and gibberellins (GAs), which all are involved in the induction of cell division or growth in microalgae **(Nguyen et al., 2020)**. Our study suggests that potential alternative strategies to counteract heavy metal toxicity could include the application of exogenous ABA or CKs (*t*Z) to improve metal uptake and algal bio-removal of heavy metals **(Figure 1)**. *Euglena gracilis* (in terms of both living and nonliving biomass) is an example of an effective microorganism for the removal of metals through its high performance as novel biosorbents **(Khatiwada et al., 2020; Moreno-Sánchez et al., 2017)**. For instance, *Euglena gracilis* was found to be tolerant to Cu, Cr, Pb, Cd, Hg and Zn **(Santiago-Martínez et al., 2015; Moreno-Sánchez et al., 2017)** and was able to highly accumulate Ni, Cd, Pb and Zn **(Sánchez-Thomas et al., 2016, 2016; García-García et al., 2018)**. This prompted researchers to investigate the metal-tolerance mechanisms of Euglena, leading to the identification of key proteins (i.e. thiol-containing proteins, sulfur assimilation pathway, transporters, stress responsive compounds, ROS detoxification, metallothionein, cell wall biosynthesis) underpinning metal tolerance and uptake (Pb, Cd, and Hg stress) via untargeted proteomics approaches **(Khatiwada et al., 2020)**. This sophisticated mechanism also was found be to regulated by CKs and ABA **(Piotrowska-Niczyporuk et al., 2020; Talarek-Karwel et al., 2020; Stirk and van Staden, 2021)**. Keeping this in mind for enhanced metal accumulation and removal, our study highlights the utility of exogenous applications of CK and ABA. Specifically, one can augment or manipulate Euglena HM removal mechanisms by phytohormones (ABA or CK) as a means to enhance HM cleanup from a contaminated environment.

### 4.2 Endogenous ABA/CK enhancement in Euglena: a new opportunity for bioremediation?

Our study found that one of the most toxic metals in nature, Pb, and two other toxic HMs, Ni and Cd, increased the level of total endogenous CKs and enhanced CK activity profiles in *E. gracilis* **(Figure 2)**. Exogenous ABA and CK improve metal uptake efficiency through the up-regulation of the CK activity profile (i.e., isoprenoid CKs, NT-CKs, MET-CKs, cZ-type CKs, iP-type CKs and total CKs all increased) **(Figure 2 and Supplementary table 2)**. This phenomenon suggests that CK profiles are responsive to HM stress, and that enhancing CK function (e.g. exogenous applied CK/over-expression of CK biosynthesis genes) might be an effective strategy to enhance accumulation and metal uptake of algal cells. The positive effect of exogenous applied CK was also found in a green alga *A. obliquus* cell under Pb stress **(Piotrowska-Niczyporuk et al., 2018, 2020; Talarek-Karwel et al., 2020)** and *Chlorella vulgaris* cells under Cd, Pb, and Cu stress **(Piotrowska-Niczyporuk et al. 2012)**. The use of exogenous CKs ameliorated the adverse effects of Ni, Pb and Cd on enhancing the pool of total endogenous CKs (2.0 fold higher (CK+Ni vs Ni); (3.3 fold higher (CK+Pb vs Pb), (1.2 fold higher (CK+Cd vs Cd)) which may indicate that an increased tendency for CK accumulation in *E. gracilis* (**Figure 2**), can mitigate metal toxicity on algal cells and improve metal uptake efficiency. Like the mode of action in algae **(Nguyen et al., 2020)**, CKs are known to activate specific signaling pathways that belong to antioxidant and phytochelatin metabolism, thereby reducing metal toxicity in higher plants - such as Cd in pea seedlings **(Al-Hakimi, 2007)**, selenium in Arabidopsis **(Jiang et al., 2019)**, arsenic in Arabidopsis **(Mohan et al., 2016)**, or Cu in tobacco (**Thomas et al., 2005)**. These coordinated activations of metal tolerance responses were also connected with aspects of our metabolomics data **(table 1d-1i)**.

Applications of exogenous ABA enhanced Pb/Cd uptake efficiency and increased the pool of endogenous CKs, which suggests there is crosstalk between ABA and CKs in *E. gracilis* during metal stress responses **(Figure 1, Figure 2 and Supplementary Table 2)**. Accordingly, the content of *c*Z- and iP-type, isoprenoid-CKs, CK-NTs, and total CKs were stimulated by ABA + Cd treatment as compared with Cd alone (**Figure 2**). Increased CK levels in algal cells treated with exogenous ABA under stress conditions may be necessary for the stabilization of the photosynthetic machinery and stimulation of cell cycle proliferation in *E. gracilis* cell exposed to metals **(Piotrowska-Niczyporuk et al., 2018)**. In summary, ABA and CK may play important roles in *E. gracilis* responses to metal stress and regulate CKs at a metabolic level, but the specific crosstalk mechanisms that modulate particular genetic signaling components remains to be determined.

### 4.3 Phytohormone crosstalk in regulating HM stress response

Cytokinins participate extensively in interactions with other hormones in microalgae under HM stress **(Piotrowska-Niczyporuk et al., 2020)**. Exogenously applied CK reduced ABA levels under metal stress (0.6 fold for CK+Ni, 0.3 fold for CK+Pb, 0.7 fold for CK+Cd, as compared to untreated controls). Thus, elevated levels of active GAs, GA_1_ and GA_3,_ (**Figure 3d**) in response to *t*Z under Ni stress conditions suggests that these growth-promoting GA forms might help modulate algal stress responses. A similar trend was found under *t*Z + Ni (4.1 fold higher for GA_1_, 28.6 fold higher for GA_3_) and *t*Z +Pb (2.4 fold higher for GA_1_, 5.7 fold higher for GA_3_) and *t*Z + Cd (2.0 fold higher for GA_1_, 3.9 fold higher for GA_3_) stresses as compared to controls (**Figure 3**). Exogenous application of ABA also increased GA_1_ (4.6 fold, p < 0.05) and GA_3_ (1.7 fold) content in *E. gracilis* cultures growing in the presence of Ni **(Figure 3)**. These results agree with data from vascular plants, indicating that exposure to heavy metals (e.g., Cd and chromium (VI)) induces expression of GA biosynthetic genes and, in turn, increases the endogenous levels of GAs which activates the specific signaling pathways and modulates gene expression involved in stress responses **(Gangwar et al., 2011; Zhu et al., 2012)**.

### 4.4 Phytohormone-modified metabolites and pathways that are associated with metal stress

Co-applications of ABA or *t*Z with metals increased the levels of metabolomic features identified as free fatty acids and those involved in lipid biosynthesis pathways (**Table 1d-1i**). One possible reason for the increase in the contents of fatty acids is that the application of ABA or CKs to algal cultures might prevent the formation of reactive oxygen species (ROS) which can attack fatty acids leading to their oxidative degradation triggered by the presence of toxic metal **(Piotrowska-Niczyporuk et al., 2012; Contreras-Pool et al., 2016; Sulochana et al., 2016)**.

The exposure of *E. gracilis* to ABA or *t*Z in combination with heavy metals resulted in significant accumulations of secondary metabolites belonging to: phenolics (m/z 473.281), terpenoids (m/z 375.178), 492.227), alkaloids (m/z 458.285, 430.334, 304.134, 580.254), flavonoids (m/z 433.077), and carotenoids (m/z 393.204). These organic compounds may play a role in metal detoxification in Euglena (**Table 1d-1i**). Bioactive compounds can act as oxygen free radical scavengers or metal chelators **(Zhai et al., 2015; Anjitha et al.,2021)**. Also, Bioactive compounds function as an upstream regulation of metallothioneins (MTs) or chelators (i.e., quercetin, catechin, anthocyanin, naringenin, puerarin), which enables a protective mechanism of phytochemicals against Cd and Pb toxicity and their physiological impacts **(Kapoor et al., 2014; Abnosi et al., 2015; Zhai et al., 2015**; **Anjitha et al.,2021; Dobrikova et al., 2021)**. Detoxification of metal ions by secondary bioactive compounds is an effective metal defense system in plants **(Lajayer et al., 2017)**. Therefore, accumulations of terpenoids, alkaloids, flavonoids, carotenoids may be correlated with the metal resistance of *E. gracilis* cultures **(Table 1d-1i)**. The roles of bioactive compounds in heavy metal chelation and tolerance (reduced oxidative-induced damages via ROS detoxification) are known from other algal species. For example, the potential antioxidant effects of nuclear-located flavonoids are of the greatest significance, as they include not only ROS quenching, but also the ability of antioxidant flavonol glycosides to chelate transition metal ions, thereby limiting the generation of ROS **(Lajayer et al., 2017; Agati et al., 2020)**. In a broad sense, this can be considered as an antioxidant function – as monoterpenes are involved in ROS detoxification in plant leaves **(Wang et al., 2017)**, and are suggested to help overcome the oxidative damages induced by high Cd and Cu levels in higher plants **(Hojati et al., 2017)**. For example, the HM hyperaccumulator *Narcissus tazetta*, when treated with Cd, was characterized by an increase in alkaloids, which have protective roles probably as ROS quenchers or for enhanced Cd accumulation **(Soleimani et al., 2020)**.

Increases in metabolite levels involved in vitamin/cofactor pathways were observed in *E. gracilis* cells treated with phytohormones. This may contribute to HM stress acclimation by scavenging free radicals and decreasing lipid peroxidation under Cd and Pb stress **(Zhai et al., 2015)**. Thus, our present data suggests that exogenously applied ABA and CK can provoke a coordinated activation of metal tolerance mechanisms leading to the increase in metal uptake efficiency which is essential for Ni, Pb and Cd sequestration.

## 5. Conclusions

Our study highlights that ABA and CKs are important regulators of algal metal accumulation/acclimation strategies based on increased metal uptake, enhanced CK metabolism, regulation of hormonal crosstalk and impacts on some core cellular metabolic pathways, all of which improve metal uptake efficiency. Finally, our results suggest that ABA and CK can form a novel strategy for metal bioremediation techniques and for sourcing microalgal value-added metabolites. As full genome sequencing, assembly, and annotation in *E. gracilis* genome has not yet been well characterized **(Ebenezer et al., 2019)**, future work into the identification of the complete genomic sequencing, pangenomic characterization of phytohormone signaling, and further comparative transcriptome analysis, and proteomic characterization in *E. gracilis* will advance insights even further.

## Abbreviations

2MeSZ: 2-methylthiol-zeatin
2MeSZR: 2-methylthiol-zeatin riboside
ABA: abscisic acid
BAP: benzylaminopurine
Cd: cadmium
*c*Z: *cis*-zeatin
*c*ZR: *cis*-zeatin 9-riboside
CK: cytokinin
DHZ: dihydrozeatin
DHZR: dihydrozeatin 9-riboside
FB: free bases (DHZ + *t*Z + cZ + iP)
GA: gibberellin
GLUC: glucosides (DZOG + *t*ZOG + *c*ZOG + DZROG + *t*ZROG
HM: Heavy metal
IAA: indole-3-acetic acid
iP: N6-isopentenyladenine
iPR: N6-isopentenyladenine 9-riboside
LC-(ESI) MS/MS: liquid chromatography-electrospray ionization-tandem mass spectrometry
Pb: lead
2MeSiP: 2-methylthiol-isopentenyladenine
2MeSiPA: 2-methylthiol-isopentenyladenosine
MET: methylthiols (2MeSZ + 2MeSiP + 2MeSZR + 2MeSiPA)
Ni: nickel
NT: nucleotides (DHZNT + *t*ZNT + *c*ZNT + iPNT)
PRM: Parallel Reaction Monitoring
RB: ribosides (DHZR + tZR + *c*ZR + iPR)
*t*Z: *trans*-zeatin
*t*ZR: *trans*-zeatin 9- riboside
XCMS: X Chromatography Mass Spectrometry

## Acknowledgment

We thank, Dr. Anna Kisiala for technical support with mass spectrometers and training with MS data analysis. We thank Noblegen Inc. for their expert advice on Euglena culture and experimental management.

## CRediT authorship contribution statement

Hai Nguyen: Conceptualization, Methodology, Validation, Formal analysis, Investigation, Writing - original draft, Visualization. Nguyen Quoc Thien: Methodology, Formal analysis, Writing - review & editing. Duc Huy Dang: Methodology, Formal analysis, Writing - review & editing. R. J. Neil Emery: Conceptualization, Methodology, Resources, Writing - original draft, Writing - review & editing, Visualization, Supervision, Project administration, Funding acquisition.

## Data availability

Data will be made available on request.

## Appendix A. Supplementary data

The following is the Supplementary files to this article:

Supplementary data

Supplementary Material and Method

