## Supplementary data for "Phytohormones enhance heavy metal responses in *Euglena gracilis*: evidence from uptake of Ni, Pb and Cd and linkages to hormonomic and metabolomic dynamics"

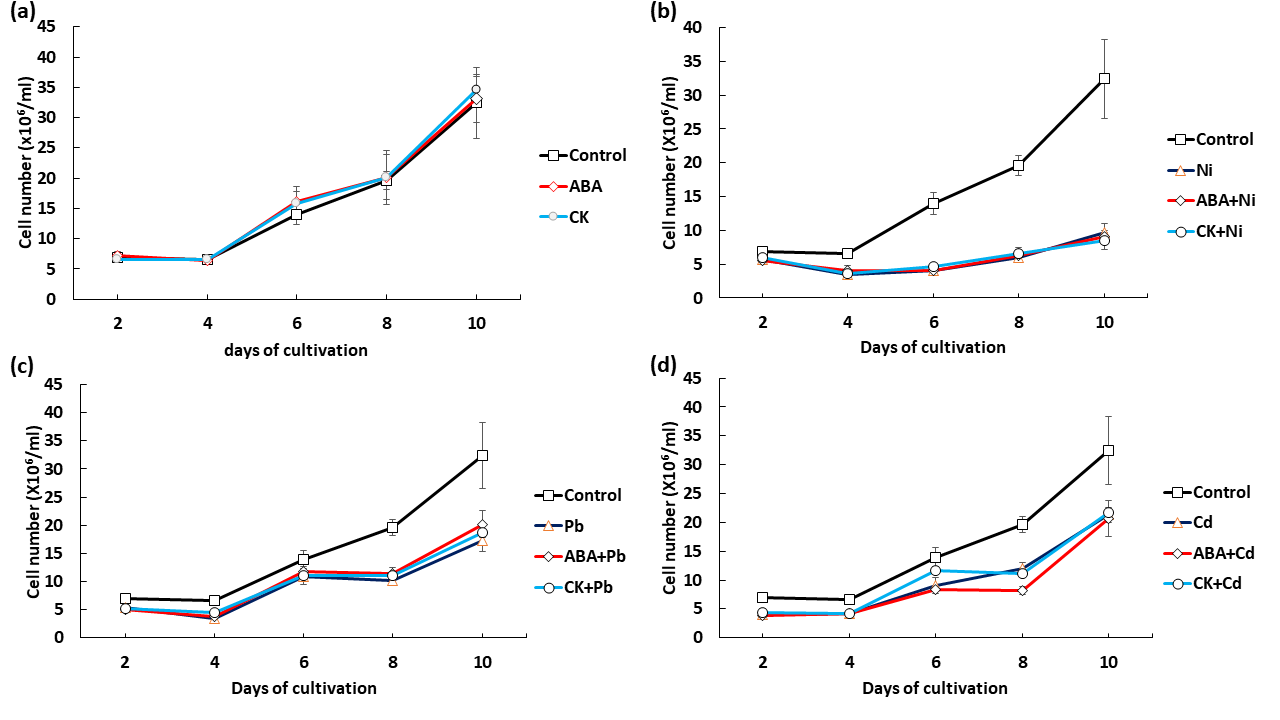
**Supplementary Figure 1**. **Phytohormones do not significantly affect cellular growth.** The effect of CK (*t*Z, 10^-9^ M) and ABA (10^-9^M) on cellular growth of *E. gracilis*. Time-course of the cellular growth of E. gracilis exposed to CK (tZ, 10^-9^ M) and ABA (10^-9^ M) and control (without treatment) (a). Time-course of the cellular growth of *E. gracilis* exposed to CK (*t*Z, 10^-9^ M) and ABA (10^-9^ M) in combination with heavy metal Nickel (0.5 mM) (b), Lead (0.1 mM) (c), Cadmium (0.025 mM) (d). Data are the means of four independent biological replications ± SD All experiments were initiated with 0.6 x 10^6^ cells/mL (t = 0) and determined by Trypan Blue staining. The difference in cell density between the treatments with phytohormone (red for ABA, green for CK) was compared to metal treatment using Dunnett's test). There were no significant differences among treatments at P> 0.05.

**Supplementary Figure 2.** (a) Bright field microscopic image (40× objective) of dead cells (dark) and living cells (green) determined by Trypan Blue staining. (b) Representative algal images after *E. gracilis* exposed to metals (Pb and Cd) and combined metal with ABA or CK (*t*Z) at the subsequent days of the treatment. Dark color indicates dead cells.

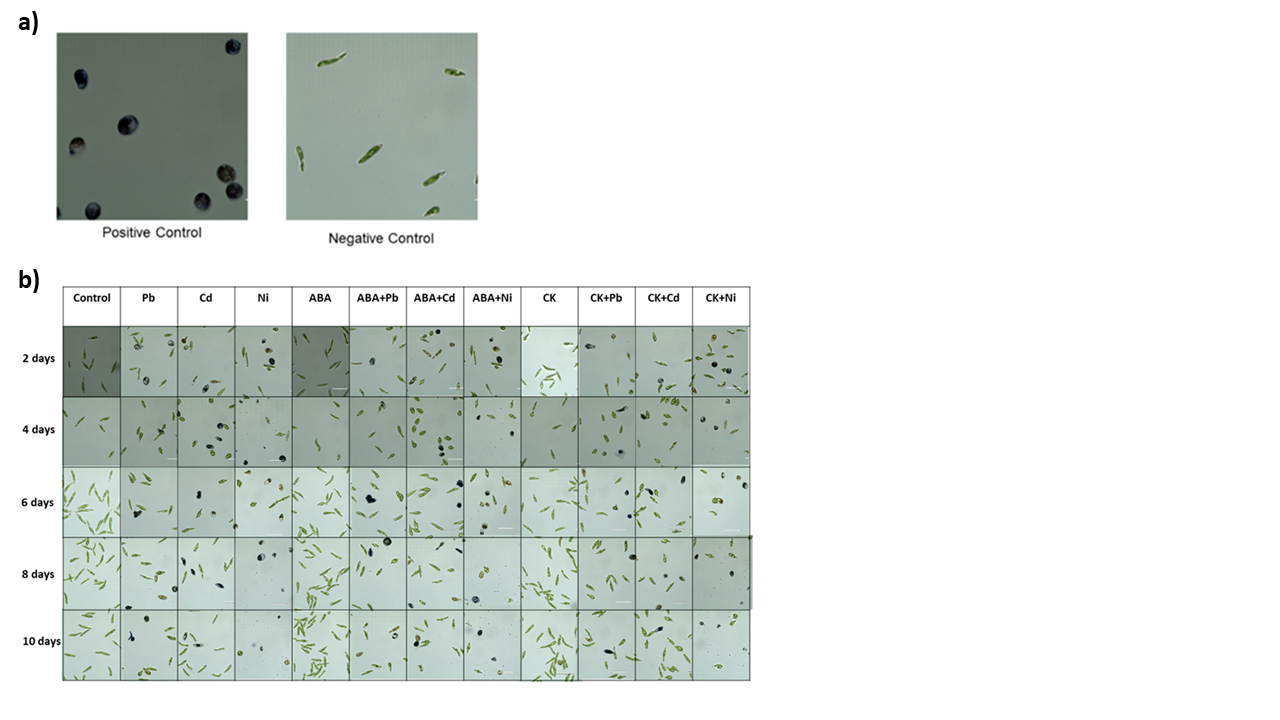

**Supplementary Table 1.**

| **Supplementary Table 1.1.** Endogenous cytokinins (CKs) and labeled CK standards scanned for by liquid chromatography-positive electrospray ionization tandem mass spectrometry (HPLC-(ESI+)-MS/MS). Labelled internal standards obtained from OlChemim Ltd. (Olomouc, Czech Republic) were used for analysis. | | |
| --- | --- | --- |
| Nucleotides (NTs) | | |
| 1. | *Trans-*zeatin riboside-5′-monophosphate (*t*ZNT) | ^2^H_5_[9RMP]Z |
| 2. | *Cis-*zeatin riboside-5′-monophosphate (*c*ZNT) |  |
| 3. | Dihydrozeatin riboside -5′-monophosphate (DHZNT) | ^2^H_3_[9RMP]DHZ |
| 4. | Isopentyladenosine-5′monophosphate (iPNT) | ^2^H_6_[9RMP]iP |
| Ribosides (RBs) | | |
| 5. | *Trans-*zeatin riboside (*t*ZR) | ^2^H_5_[9R]Z |
| 6. | *Cis-*zeatin riboside (*c*ZR) |  |
| 7. | Dihydrozeatin riboside (DHZR) | ^2^H_3_[9R]DHZ |
| 8. | Isopentyladenosine (iPR) | ^2^H_6_[9R]iP |
| Free bases (FBs) | | |
| 9. | *Trans-*zeatin (*t*Z) | ^2^H_3_DHZ |
| 10. | *Cis-*zeatin (*c*Z) |  |
| 11. | Dihydrozeatin (DHZ) |  |
| 12. | Isopentyladenine (iP) | ^2^H_6_iP |
| Glucosides (GLUCs) | | |
| 13. | *Trans*-zeatin-O-glucoside (*t*ZOG) | ^2^H_5_ZOG |
| 14. | *Cis*-zeatin-O-glucoside (*c*ZOG) |  |
| 15. | Dihydrozeatin-O-glucoside (DHZOG) | ^2^H_7_DHZOG |
| 16. | *Trans*-zeatin-O-glucoside riboside (*t*ZROG) | ^2^H_5_ZROG |
| 17. | *Cis*-zeatin-O-glucoside riboside *c*ZROG |  |
| 18. | Dihydrozeatin-O-glucoside riboside (DHZROG) | ^2^H_7_DHZROG |
| 19. | *Trans*-zeatin-9-glucoside (*t*Z9G) | ^2^H_5_Z9G |
| 20. | *Cis*-zeatin-9-glucoside (*c*Z9G) |  |
| 21. | Dihydrozeatin-9-glucoside (DHZ9G) | ^2^H_3_DHZ9G |
| 22. | Isopentenyladenine-7-glucoside (iP7G) | ^2^H_5_iP7G |
| 23. | Isopentenyladenine-9-glucoside (iP9G) |  |
| Methylthiols (METs) | | |
| 24. | 2-Methylthio-*trans*-zeatin (2MeSZ) | ^2^H_5_MeSZ |
| 25. | 2-Methylthio-*trans*-zeatin riboside (2MeSZR) | ^2^H_5_MeSZR |
| 26. | 2-Methylthio-isopentyladenine (2MeSiP) | ^2^H_6_MeSiP |
| 27. | 2-Methylthio-isopentyladenosine (2MeSiPA) | ^2^H_6_MeSiPR |
| Aromatic Cytokinins | | |
| 28. | Benzylaminopurine (BA) | ^2^H_7_BA |
| 29. | Benzylaminopurine riboside (BAR) | ^2^H_7_BAR |
| **Supplementary Table 1.2**. Names and abbreviations of endogenous phytohormone and labeled phytohormone standards scanned for by liquid chromatography-negative electrospray ionization tandem mass spectrometry (HPLC-(ESI-)-MS/MS). Deuterated internal standards purchased from OlChemim Ltd. (Olomouc, Czech Republic) were used to identify and quantify phytohotmone. | | |
| 30 | ABA | ^2^H_6_ABA |
| 31 | GA1 | ^2^H_2_GA1 |
| 32 | GA3 | ^2^H_2_GA3 |
| 33 | GA4 | ^2^H_2_GA4 |
| 34 | GA7 | ^2^H_2_GA7 |
| 35 | IAA | ^2^H_5_IAA |
| 36 | SA | ^2^H_6_SA |

**Supplementary Table 2.** The effect of exogenous ABA and cytokinin (*t*Z) on the level of different forms of endogenous CKs in *E. gracilis* cells exposed to metals (nickel, lead, and cadmium) and ABA or CK (*t*Z) on the 10th day of cultivation in relation to control cells. Data are the means of four independent biological replications (n= 4 ± SE). Asterisks denote significant differences among treatments in a phytohormone level compared to the control culture (ANOVA; Dunnett's test, P < 0.05). The lack of asterisks indicates there were no significant differences. nd, not detected.

| **CK content [pmol/gDW]** | | **Treatment** | | | | | | | | | |
| --- | --- | --- | --- | --- | --- | --- | --- | --- | --- | --- | --- |
|  |  | **Control** | **Ni** | **Pb** | **Cd** | **ABA + Ni** | **ABA + Pb** | **ABA + Cd** | **CK + Ni** | **CK + Pb** | **CK + Cd** |
| **Free bases** | **tZ** | nd | nd | nd | nd | nd | nd | nd | nd | nd | nd |
|  | **cZ** | nd | nd | nd | nd | nd | nd | nd | nd | nd | nd |
|  | **DHZ** | nd | nd | nd | nd | nd | nd | nd | nd | nd | nd |
|  | **iP** | nd | nd | nd | nd | nd | nd | nd | nd | nd | nd |
| **Ribosides** | **tZR** | nd | nd | nd | 9.23±  3.51 | nd | nd | nd | nd | nd | nd |
|  | **cZR** | nd | nd | nd | nd | nd | nd | nd | nd | nd | nd |
|  | **DHZR** | nd | nd | nd | nd | nd | nd | nd | nd | nd | nd |
|  | **iPR** | nd | nd | nd | 1.95±  0.77 | 61.29±  43.35 | nd | nd | nd | nd | nd |
| **Nucleotides** | **tZNT** | nd | nd | nd | nd | nd | nd | nd | nd | nd | nd |
|  | **cZNT** | 27.68± 5.11 | 65.49± 5.39 | 20.51± 2.00 | 45.30± 5.60 | 40.19± 17.22 | 29.88± 13.74 | 98.79± 16.02 | 83.47± 27.87 | 26.70± 5.88 | 66.27± 9.98 |
|  | **DHZNT** | nd | nd | nd | nd | nd | nd | nd | nd | nd | nd |
|  | **iPNT** | 240.20± 88.94 | 848.06± 116.61 | 145.45± 18.41 | 764.13± 172.30 | 565.53± 177.60 | 257.65± 43.22 | 3258.67± 503.76 | 1738.10± 578.39 | 531.00± 226.76 | 890.79± 432.98 |
| **Methylthiols** | **2MeSZ** | 2.53±  0.85 | 4.29±  1.22 | 3.40±  1.23 | 3.82±  1.61 | nd | nd | 0.66±  0.55 | 10.87±  7.88* | nd | nd |
|  | **2MeSZR** | nd | nd | 1.74±  0.65 | 1.53±  0.43 | nd | nd | 1.56±  0.65 | nd | nd | nd |
|  | **MeSiP** | nd | nd | nd | nd | nd | nd | nd | nd | nd | nd |
|  | **MeSiPA** | nd | nd | nd | nd | nd | nd | nd | nd | nd | nd |
| **Aromatic CK** | **oT** | nd | nd | nd | 0.08±  0.03 | nd | 0.39±  0.16* | 0.05±  0.02 | nd | nd | nd |
|  | **BAP** | 1.50±  0.57 | 2.28±  0.56 | 1.80±  0.33 | 0.70±  0.14 | 8.12±  8.25* | 1.53±  0.44 | 0.37±  0.10 | 4.19±  0.94 | 4.96±  4.33 | 7.66±  3.37 |
| **Total CK** |  | 271.05± 82.56 | 919.16± 123.60 | 172.40± 19.31 | 825.48± 175.26 | 681.42± 127.80 | 289.26± 52.92 | 3359.99± 520.56 | 1842.33± 397.51 | 566.40± 197.67 | 964.94± 312.18 |

| **Supplementary Table 3.** The effect of CK (tZ, 10^-9^ M) and ABA (10^-9^M) on cellular growth of E. gracilis. Data are the means of four independent biological replications (n= 4 ± SE). | | | | | | | | | | | |
| --- | --- | --- | --- | --- | --- | --- | --- | --- | --- | --- | --- |
| **Cell number (x10^6^/ml)** | | **Treatment** | | | | | | | | | |
|  |  | **Control** | **Ni** | **Pb** | **Cd** | **ABA + Ni** | **ABA + Pb** | **ABA + Cd** | **CK + Ni** | **CK + Pb** | **CK + Cd** |
| **Day 2** | Mean±SD | 6.895±  0.775 | 5.753±  0.313 | 5.315±  0.599 | 3.985±  0.740 | 5.505±  0.334 | 5.085±  0.385 | 3.855±  0.114 | 6.015±  0.996 | 5.1400±  0.370 | 4.42±  0.657 |
|  | N | 4 | 4 | 4 | 4 | 4 | 4 | 4 | 4 | 4 | 4 |
| **Day 4** | Mean | 6.595±  0.507 | 3.400±  0.404 | 3.485±  0.295 | 4.13±  0.380 | 4.0700±  0.688 | 3.685±  0.115 | 4.1900±  0.320 | 3.6800±  0.301 | 4.4200±  0.239 | 4.15±  0.221 |
|  | N | 4 | 4 | 4 | 4 | 4 | 4 | 4 | 4 | 4 | 4 |
| **Day 6** | Mean | 13.935±  1.657 | 4.1025±  0.348 | 10.8500±  0.909 | 9.03±  1.331 | 3.9900±  0.657 | 11.79± 1.159 | 8.365± 0.237 | 4.673± 0.354 | 11.100± 1.615 | 11.73± 0.650 |
|  | N | 4 | 4 | 4 | 4 | 4 | 4 | 4 | 4 | 4 | 4 |
| **Day 8** | Mean | 19.620± 1.458 | 5.925±  0.355 | 10.230±  0.544 | 11.930±  1.086 | 6.24±  0.935 | 11.410±  1.042 | 8.150±  0.670 | 6.615±  0.792 | 11.060±  0.594 | 11.08±  0.444 |
|  | N | 4 | 4 | 4 | 4 | 4 | 4 | 4 | 4 | 4 | 4 |
| **Day 10** | Mean | 32.425±  5.844 | 9.615±  1.335 | 17.225±  1.898 | 21.275±  1.345 | 9.090±  0.866 | 20.15±  2.390 | 20.64±  3.125 | 8.46±  1.293 | 18.625±  1.670 | 21.62±  0.389 |
|  | N | 4 | 4 | 4 | 4 | 4 | 4 | 4 | 4 | 4 | 4 |

| **Supplementary Table 4.** The effect of CK and ABA on removal capacity (%) of E. gracilis for three heavy metals (Lead (0.1 mM), Cadmium (0.025 mM), Nickel (0.5 mM)). Data are the means of at least three independent biological replications ± SD. | | | | | | | | | | |
| --- | --- | --- | --- | --- | --- | --- | --- | --- | --- | --- |
| **Metal removed/culture (%)** | | **Treatment** | | | | | | | | |
|  |  | **Ni** | **Pb** | **Cd** | **ABA + Ni** | **ABA + Pb** | **ABA + Cd** | **CK + Ni** | **CK + Pb** | **CK + Cd** |
| **Day 2** | Mean±SD | 73.087±  0.443 | 4.987±  0.604 | 27.487±  0.939 | 74.064±  0.138 | 8.601±  0.583 | 31.908±  0.371 | 74.056±  0.062 | 8.619±  1.325 | 32.840±  0.704 |
|  | N | 4 | 4 | 4 | 4 | 4 | 4 | 4 | 4 | 4 |
| **Day 4** | Mean | 73.632±  0.299 | 13.306±  0.737 | 48.213±  3.170 | 73.875±  0.131 | 22.020±  4.263 | 40.241±  1.008 | 73.283±  0.223 | 16.300±  1.394 | 39.118±  1.329 |
|  | N | 4 | 4 | 4 | 4 | 4 | 4 | 4 | 4 | 4 |
| **Day 6** | Mean | 76.750±  2.409 | 4.846±  0.970 | 41.969±  0.350 | 75.249±  1.195 | 7.139± 1.148 | 45.753± 0.490 | 74.035± 0.058 | 8.900± 1.283 | 48.191±  1.197 |
|  | N | 4 | 4 | 4 | 4 | 4 | 4 | 4 | 4 | 4 |
| **Day 8** | Mean | 73.688±  1.372 | 6.703±  2.370 | 47.598±  1.330 | 72.913±  0.307 | 9.914±  5.580 | 58.152±  5.705 | 73.913±  0.571 | 6.351±  1.058 | 46.221±  0.713 |
|  | N | 4 | 3 | 4 | 4 | 4 | 4 | 4 | 4 | 4 |
| **Day 10** | Mean | 70.079±  0.292 | 0.850±  0.438 | 52.304±  2.749 | 72.830±  2.283 | 5.273±  0.318 | 56.572±  3.450 | 71.214± 0.527 | 10.204±  2.952 | 61.518±  1.711 |
|  | N | 4 | 3 | 4 | 4 | 3 | 4 | 4 | 3 | 4 |

**Supplementary table 5.** Abundance of metabolites identified in *E. gracilis* under different treatments. Biological triplicates were used for all experiments.

| **(FC) Fold change ±1.5, t-test p-value < 0.05.** | | | | |
| --- | --- | --- | --- | --- |
| **Treatments** | **Initially indentified features (before extracted ion chromatograms selection)** | **Metabolites changed in relative abundance⁎** | **Metabolites increased in abundance (>1.5 FC)** | **Metabolites decreased in abundance (<1.5 FC)** |
| **Ni vs Control** | 3108 | 13 | 1 | 12 |
| **Pb vs Control** | 2646 | 16 | 4 | 12 |
| **Cd vs Control** | 2670 | 12 | 6 | 6 |
| **(ABA + Ni) vs Ni** | 2919 | 7 | 5 | 2 |
| **(ABA + Pb) vs Pb** | 2866 | 6 | 1 | 5 |
| **(ABA + Cd) vs Cd** | 3055 | 3 | 2 | 1 |
| **(CK + Ni) vs Ni** | 2225 | 6 | 4 | 2 |
| **(CK + Pb) vs Pb** | 2156 | 7 | 2 | 5 |
| **(CK + Cd) vs Cd** | 1900 | 1 | 1 | 0 |

**Supplementary Materials and Methods**

**2.1 Euglena growth conditions and treatments**

Axenic cultures of *Euglena gracilis* Klebs (CPCC 469) were purchased from Canadian phycological culture centre, department of Biology, University of Waterloo. Cultures were grown axenically in 50 mL of Modified Acid Medium (MAM) (**Olaueson and** **Stokes, 1989**), at pH 4.5, in 250 mL plastic flasks. Euglena cultures were maintained on a shaker in an environmental chamber at 24°C. Lighting was supplied by cool-white fluorescent light with an intensity of 97.3 Par under 12/12 light/dark cycle until culture reached 10^6^ cell/mL.

**2.2 Determination of cell growth**

The concentrations of applied compounds (heavy metals and phytohormones) were selected based on growth-curve trial experiments. The heavy metals tested against *E. gracilis* were Pb (0.1 mM and 0.2 mM Pb), Ni (0.5 mM, 1 mM, and 2 mM Ni) and Cd (1.5 × 10^-3^ mM, 0.025 mM, and 0.05 mM Cd) to determine a minimum inhibitory concentration (at least 35% inhibition but without prevailing mortality) **(Sánchez-Thomas etal., 2016; Khatiwada et al., 2020)**. Hormone concentrations were tested (10^-3^ M, 10^-6^ M, 10^-9^ M for ABA and CK) to determine which were the most effective for inducing of cell proliferation. The selected metal and hormone concentrations were used for subsequent experiments (part 2.4, 2.5 and 2.6). Briefly, phytohormones ABA (10^-9^ M) and CK (10^-9^ M *t*Z) were used on their own and in combination with 0.1 mM Pb (as Pb(NO_3_)_2_), 0.05 mM Ni (as NiCl_2_.6H_2_O), or 0.025 mM Cd (as Cd(NO_3_)_2_.4H_2_O). In all testing conditions, there was a control (non-treatment) and nine treatments (Ni; Pb; Cd; ABA; *t*Z; ABA+Ni, ABA+Pb, ABA+Cd, *t*Z+Ni, *t*Z+Pb, *t*Z+Cd). Exogenous phytohormones and metals used in experiments were purchased from OlChemim Ltd (Olomouc, Czech Republic) and Thermo Fisher Scientific, respectively. All supplements were added to 50 mL cultures (using MAM medium) during the early exponential phase (Day 0 with 0.6 x 10^6^ cells/mL) with 4 biological replicates (4 separate 250 mL flasks).

For cellular growth characterization of *E. gracilis*, 1 mL samples were collected from treatments after 2, 4, 6, 8 and 10 days of growth and the number of cells was determined by Trypan Blue staining **(Khatiwada et al., 2020)**. Dead cells stained blue as Trypan Blue solution diffused in and stained the cells whereas viable cells remained green, without the penetration of the stain. Enumeration of living Euglena cells was determined by counting cells immobilized with 0.05% HCl, using a 0.1 mm Neubauer hemocytometer (Thermo Fisher Scientific, USA) and a light microscope (Life Technologies AMAFD1000) coupled with EVOS cell imaging systems (Thermo Fisher Scientific, California, USA). Collected cells at the end of treatments (10 days) were washed by deionized water and freeze-dried using a FreeZone freeze dryer (Labconco, Missouri, USA) for subsequent analysis. Details of the cell growth are described in **Supplementary Figure 1,2 and supplementary table 3**.

**2.3 Determination of metal accumulation by Euglena using ICP-MS**

At the end of each exposure time, 1mL of sample was filtered using a 0.2 μm syringe tip filter (Canadian Life Science), acidified with double-distilled trace metal grade HNO_3_, and stored at room temperature until elemental analysis. Metal concentrations were determined utilizing a triple quadrupole inductively coupled plasma mass spectrometer (ICP-MS, 8800 Agilent Technologies) at the Water Quality Centre, Trent University following the protocol previously described by **Dang et al., (2020**). Briefly, the instrument was operated under He mode to remove polyatomic interferences and In was used as an internal standard at a final concentration of 20 µg L^-1^. Analytical recovery was determined using a certified reference material (ES-L1, SCP Science, Quebec, Canada). We also determine metal concentrations in the stock metal solution, blank sample (medium only), Euglena sample (Euglena only), and phytohormone samples (Euglena with phytohormone) for blank and control conditions.

The uptake of metals by *Euglena* was computed using changes in the metal concentration in the test medium during the exposure period expressed as percentage removal: $\frac{R_{0}-R_{i}}{R_{0}}\times100$, where: R_0_ - initial concentration-day 0 and R_i_ - concentration of metal after each exposure period **(Kong et al., 2019; Zada et al., 2021).** More than three biological replicates were used for the ICP-MS analysis **(Supplementary Table 4)**.

**2.4 Cytokinin metabolite analysis**

**2.4.1 Cytokinin extraction**

Algal cell pellets (25 mg dry weight) were harvested at the end of the experiment as described above and four replications at final time point (10 days) were used to determine endogenous cytokinin (CK) levels. Extraction, purification, and quantification of CKs were performed as previously described (**Kisiala et al., 2019)**. Samples were extracted in 1 mL of Bieleski solution to which 29 stable isotope-labelled internal standards were added (20 ng of each of the deuterated internal standards, OlChemim Ltd. (Olomouc, Czech Republic)) corresponding to the different forms of CK that were scanned for as in previous studies **(Supplementary Table 1)** **(Kisiala et al., 2019; Aoki et al., 2021; Bean et al.,2021; Palberg et al., 2022)**. Chilled samples were homogenized using a ball mill (Retsch MM300, Qiagen, Valencia, CA, USA) and allowed to passively extract overnight at −20 °C in modified Bieleski extraction solvent. Samples were centrifuged (10,000×g for 10 min), and the supernatant collected. The remaining pellets were re-suspended in an additional 1 mL of the extraction solvent and allowed to extract for an additional 30 min at −20 °C. Re-extracted samples were centrifuged and both supernatants were pooled and evaporated to dryness in a speed vacuum concentrator (Savant SPD111V, UVS400, Thermo Fisher Scientific, Waltham, MA, USA) at room temperature. Dried residues were reconstituted in 1 mL of 1 M HCO_2_H to ensure complete protonation of all CKs. Each extract was subjected to solid phase extraction (SPE) on a mixed mode, reversed-phase/cation-exchange cartridges (MCX 6cc; 200 mg, Canadian Life Sciences, Peterborough, ON, Canada). Cartridges were activated with methanol (CH_3_OH) and equilibrated with 1M formic acid HCO_2_H. Each sample was loaded and allowed to pass through the column by gravity and the columns washed with 1 M HCO_2_H, followed by CH_3_OH. The nucleotide CKs were eluted first using NH_4_OH. Other CK groups were retained on the column based on charge and their hydrophobic properties. Free bases, ribosides, methylthiols, and glucosides were subsequently eluted together using 0.35 M NH_4_OH in 60% CH_3_OH. The nucleotide samples were dephosphorylated using three units of alkaline phosphatase in 1 mL of 0.1 M ethanolamine-HCl (pH 10.4) for 12 h at 37 °C. Both eluted CK fractions were evaporated in a speed vacuum concentrator and stored at −20 °C until further processing. Prior to UHPLC-(ESI+)-MS/MS analysis, the CKs were re-constituted in initial mobile phase conditions (95:5 H_2_O:CH_3_CN) with 0.08% CH_3_CO_2_H. Samples were transferred to glass auto-sampler vials (Agilent, United States) and stored at −20 °C until analysis.

**2.4.2 CK Quantification using UHPLC-(ESI+)-HRMS/MS**

Endogenous CKs within *E. gracilis* samples were identified and quantified using UHPLC-(ESI+)-HRMS/MS (**Kisiala et al., 2019**). A 25 µl sample volume was injected into the Thermo Ultimate 3000 UHPLC coupled to a Thermo Q-Exactive™ Orbitrap mass spectrometer equipped with a heated electrospray ionization (HESI) source (Thermo Scientific, San Jose, CA, USA). Compounds were separated using a reversed-phase C18 column (Kinetex 2.6 µ C18 100 A, 2.1 × 50 mm; Phenomenex, Torrance, CA, USA). All hormone forms were eluted with a multistep gradient of component A: H_2_O with 0.08% CH_3_CO_2_H mixed with component B: CH_3_CN with 0.08% CH_3_CO_2_H at a flow rate of 0.5 mL/min. The initial conditions of 5% B were held for 0.5 min, increasing linearly to 45% B over 4.5 min followed by an increase to 95% B over 0.1 min; and held at 95% B for 1 min, before returning to initial conditions for 2 min of column re-equilibration, total run time was 8.2 min.

The eluate was introduced into the Orbitrap HESI source (capillary temperature of 250°C) and analyzed using parallel reaction monitoring (PRM) at 35,000 resolution. CKs were analyzed in positive ion mode. The HESI source was operated with sheath gas, 30 arbitrary units; auxiliary gas, 8 arbitrary units; max spray current, 100 µA; auxiliary gas heater temperature, 450 °C; S-lens RF level, 60 and spray voltage 3.9 kV. The PRM parameters included the following: automatic gain control (AGC), 1 × 10^6^; maximum injection time, 128 ms; m/z 1.2 isolation window and normalized collision energy individually optimized for each CK analyte.

All data were analysed using Thermo Xcalibur (v. 4.1) software (Thermo Scientific, San Jose, CA, USA), to calculate peak areas. Quantification was achieved through isotope dilution analysis based on recovery of ^2^H-labelled internal standards. Blank media samples (control, MAM solution) were processed and analysed using the same methodologies shown no background levels of CKs. Four biological replicates were used for the cytokinin metabolite analysis **(Supplementary Table 2)**.

**2.5 Hormonomics analysis**

ABA, indoleacetic acid (IAA), SA, and selected Gibberellins (GA_1_, GA_3_, GA_4_, GA_7_) were scanned for in 25 mg dry weight *E. gracilis* pellet samples collected after 10 days of treatment. Extraction, purification, and quantification of phytohormones was performed as described by **Cao et al (2020)** with modifications. Samples were extracted in 1 mL of 50% acetonitrile (CH_3_CN) solution and spiked with 10 ng stable isotope-labelled internal standards (Sigma-Aldrich, Inc., St. Louis, MO, USA). Pellets were removed by centrifugation (10 min at 10,000 g), and the supernatants collected, and pellets were re-extracted with 0.5 mL of ice-cold 50% CH_3_CN solvent for 30 min at 4°C. The solids were separated by centrifugation and discarded, and the pooled supernatants were evaporated to dryness at 35°C under vacuum (Model SPD111V; ThermoFisher Scientific, Ottawa, Canada). The extraction residues were reconstituted in 3 mL of 50% CH_3_CN. The hormone-containing extracts were purified and concentrated on HLB cartridges (Canadian Life Sciences, Peterborough, ON, Canada). Cartridges were activated with HPLC grade CH_3_OH and equilibrated using 2 mL of 30% CH_3_CN. After cartridge equilibration, each sample was loaded with 300 µL 30% CH_3_CN, spun down at 3000 rpm and transferred to glass auto-sampler vials (Agilent, United States) and stored at −20 °C.

All samples were analyzed in negative ion mode using a HPLC–ESI–MS/MS (QTrap5500 triple quadrupole mass spectrometer (Sciex Applied Biosystems, Massachusetts, United States) connected to a Shimadzu LC10ADvp equipped with PE200 autosampler. A 20 uL aliquot was injected onto a reverse phase C18 column (Kinetex 2.6u C18 100 A, 2.1 × 50 mm; Phenomenex). Phytohormones (ABA, IAA, GA_1_, GA_3_, GA_4_, GA_7_ and SA) were eluted using component A: H_2_O with 0.5% formic acid (FA) and component B: 0.5% FA in ACN (v/v), at a flow rate of 0.5 mL min^−1^. Phytohormone was analyzed in negative ionization mode and the programmed step gradient was followed by **Cao et al., 2020**. Hormone concentrations were quantified based on the peak area ratios of endogenous to corresponding recovered internal standard **(Cao et al., 2020).** Raw phytohormone data were analyzed with Analyst 1.6.2 software (AB SCIEX, Concord, ON, Canada). The levels of ABA, IAA, GA_1_, GA_3_, GA_4_, GA_7_ and SA are reported as the average of 4 replicates (pmol/g dwt). Blank media samples (control, MAM solution) were analysed using the same methodologies shown no matrix effect of phytohormone.

**2.6 High resolution Orbitrap mass spectrometry-based metabolomics**

For the metabolomics extractions, 25 mg dry weight pellet samples were collected from experiments described above and re-suspended in 1 mL of ice-cold 80% MeOH, sonicated for 15 min, and centrifuged at 10,000 rpm for 10 min. Next, 700 µL of supernatant were transferred to 0.2 µM PVDF centrifuge filters with a 2 mL receiver tube and centrifuged at 4000 rpm for 3 min. Filtered extracts were dried down at ambient temperature in a speed vacuum concentrator. Residues were re-dissolved in 400 µL 50% MeOH/Water and spun down again at 10000 rpm for 10 min **Renaud et al. (2017)**. Subsequently, 200 µL of supernatant were transferred to a 400 µL auto sampler inserts inside a 2 mL glass vial (Agilent, United States) with septa cap. Labeled standards (10 ng of [^2^H_7_]BA and 10 ng of [^2^H_6_]ABA) were added to use as a lock mass and monitor retention times across samples. Full scan analyses were performed following **Renaud et al. (2017)** with modifications, using UHPLC-MS. The samples were passed into the mass spectrometer from a Thermo Ultimate 3000 UHPLC system ((FisherScientific, Ottawa, Canada).). For full scan analysis, each sample was run in negative ion mode over the mass range of m/z 80−600, at 35,000 resolutions, with automatic gain control (AGC) target of 3 × 10^6^, and maximum injection time (IT) of 128 ms. For metabolite selection, data files (as Thermo.RAW) were converted to mzXML file by ProteoWizard (Version 2.1) and uploaded to XCMS Online (https://xcmsonline.scripps.edu) to query significant differences among the metabolite profiles **(Papaioannou et al., 2021; Rodríguez-Moro et al., 2022)**. Parameters for the analysis were selected as outlined by **Chen et al. (2017)**, with modifications to tailor the method to our UHPLC. Feature detection was completed with the centWave method and a ppm tolerance of 5. The pre-filter was set to 6 scans with a minimum 5,000 intensity, the signal to noise threshold was 5 and noise was set to 1× 10^6^ for use in negative mode. Retention time correction was conducted using the obiwarp method; the grouping included features present in at least 25% of all samples, the allowable retention time deviation was 10 s, and the m/z width was set to 0.01. Criteria for feature selection included: retention time greater than 1.5 min and less than 6 min to avoid solvent fronts and tails during the pressure rebound of the HPLC; a fold change greater than 1.5; and a p < 0.05. A visual representation of peaks in XCMS Online was used to check peak shape and intensity that had to be significantly greater than baseline noise. To assure the sensitivity, reliability and validity of metabolite profiling analysis, specific internal standards were monitored in both negative mode (ABA) and positive mode (Benzylaminopurine), and samples were analyzed at three dilution levels (1x, 10x and 100x dilution factor) (**Aoki et al., 2019; Palberg et al., 2022)**. The optimization tests showed that undiluted samples in negative mode were the most robust and so these parameters were selected for subsequent analysis. High abundance metabolites (fold change ±1.5, p-value < 0.05) in the *E. gracilis* are summaried in **Figures 4**. Supervised hierarchical clustered heatmaps showed clear differences between the different treatment groups and a high degree consistency of biological replication within each treatment **(data not shown)**. High abundance metabolites (fold change ±1.5, p-value < 0.05) in the *E. gracilis* according to the High Resolution Orbitrap Mass Spectrometry-based metabolomics analysis are summarized in **Supplementary table 5** and **Figure 4**. Examples of highly selective extracted ion chromatograms (EIC of significantly up/down regulated metabolites are shown in **Supplementary Figures 3.1- 3.9**.

**Metabolite Identification**

All putative feature identifications corresponded to a Level 3 confidence assignments – according to a scale set by the Chemical Analysis Working Group (CAWG) of the Metabolomics Standards Initiative (MSI) and as modifed by **Schrimpe-Rutledge, 2016**. This means that accurate masses of features were queried against databases to produce tentative structures and identifications.

MetaboQuest (http://tools.omicscraft.com/MetaboQuest/) was used as a single search tool and was queried with the m/z value of each unfragmented feature as a positive ion, with [M+H], [M+Na], [M+ACN] adducts specified. MetaboQuest provided suggestions for identities to be confirmed by cross-referencing with the precursor ion m/z values in databases such as KEGG (<https://www.genome.jp/kegg/compound/>) and METLIN (http://metlin.scripps.edu). Functional annotations of the proposed metabolites were derived using MetaboQuest (<http://tools.omicscraft.com/MetaboQuest/>), PubChem (https://pubchem.ncbi.nlm.nih.gov/) and KEGG **(Cheng et al., 2022 Rodríguez-Moro et al., 2022)**. Based on functional annotation, only the high abundance metabolites that may play a role in heavy metal tolerance and accumulation were categorized and shown**.**

**Statistical analysis**

Statistical analyses were carried out by SPSS Software (IBM, version 28.0) on algal physiology and biochemistry data using a one-way analysis of variance (ANOVA) followed by Dunnett's post-hoc test to test for significant effects of treatments when compared to the respective control. Significant differences refer to a P-value of < 0.05 for the Dunnett's test are refer to significant effects of treatments as compared to control (non-treated group). Data points and error bars reflect means ± standard errors of 4 biological replicates for phenotyping and phytohormone data and at least three biological replicates for metal uptake analyses.
